# Structural basis of client specificity in mitochondrial membrane-protein chaperones

**DOI:** 10.1101/2020.06.08.140772

**Authors:** Iva Sučec, Yong Wang, Ons Dakhlaoui, Katharina Weinhäupl, Tobias Jores, Doriane Costa, Audrey Hessel, Martha Brennich, Doron Rapaport, Kresten Lindorff-Larsen, Beate Bersch, Paul Schanda

## Abstract

Chaperones are essential for assisting protein folding, and for transferring poorly soluble proteins to their functional locations within cells. Hydrophobic interactions drive promiscuous chaperone–client binding, but our understanding how additional interactions enable client specificity is sparse. Here we decipher what determines binding of two chaperones (TIM8·13, TIM9·10) to different integral membrane proteins, the all-transmembrane mitochondrial carrier Ggc1, and Tim23 which has an additional disordered hydrophilic domain. Combining NMR, SAXS and molecular dynamics simulations, we determine the structures of Tim23/TIM8·13 and Tim23/TIM9·10 complexes. TIM8·13 uses transient salt bridges to interact with the hydrophilic part of its client, but its interactions to the trans-membrane part are weaker than in TIM9·10. Consequently, TIM9·10 is outcompeting TIM8·13 in binding hydrophobic clients, while TIM8·13 is tuned to few clients with both hydrophilic and hydrophobic parts. Our study exemplifies how chaperones fine-tune the balance of promiscuity *vs*. specificity.

## Introduction

Cellular survival and function fundamentally rely on an intact proteome. Proteins within cells need to be correctly folded to their functional conformation, and be present at the cellular location of their function. Chaperones play a central role in maintaining this cellular protein homeostasis (1), by either helping other proteins to reach their functional threedimensional structure after synthesis, by transporting them across the cytosol or organelles, or by sustaining their native fold along their lifetime. More than 20,000 proteins are required to fulfill the functions of human cells, and it is believed that the majority relies on chaperones to reach and maintain their native fold (2). Given the diversity of the client proteins, many chaperones promiscuously interact with tens of different ‘client’ proteins that may differ widely in size, structure and physico-chemical properties. However, the need for efficient binding and refolding of their clients also calls for some degree of specificity. Chaperones operate at this delicate balance of promiscuity and specificity to their clients. The interactions determining the chaperone–client specificity are only partly understood.

Hydrophobic interactions play a crucial role for chaperone interactions as most chaperones bind to hydrophobic patches on their clients and shield them from aggregation. Electrostatic charges also play a role in some chaperone complexes (3). The exact nature of the interaction motifs recognized by different chaperones differ (4). For example, the interaction with the Hsp70 chaperone family is particularly dependent on the presence of Ile, Phe, Leu and Val (5, 6); the SecB chaperone recognizes 9-residue long stretches enriched in aromatic and basic residues (7); the chaperone Spy uses longer-range charge interactions for the formation of an initial encounter complex, followed by more tight binding mediated by hydrophobic interactions, (8) whereby structurally frustrated sites on the client protein are particularly prone to binding (9).

Our understanding of the underlying principles of chaperone-client interactions is hampered by the lack of atomic-level structural views and dynamics of these complexes. Their inherently dynamic and often transient nature represent significant experimental challenges towards structural characterization. Only a very limited number of chaperone complex structures have been reported (reviewed in (10)). The modes of interactions that they revealed range from rather welldefined binding poses of client polypeptides in the chaperone’s binding pockets, reminiscent of classical protein complexes, to highly flexible ensembles of conformations (’fuzzy complexes’). In the latter, a multitude of local chaperone-client interactions may result in a high overall affinity despite the low affinity and short life time of each individual intermolecular contact.

Multiple molecular chaperones are present in the cell with mutually overlapping functions and ‘clientomes’ (2, 11, 12). It is poorly understood, however, whether a given client protein adopts a different conformation (or ensemble of conformations) when it is bound to different chaperones, and if different clients, when bound to a given chaperone, all show similar conformational properties. α-Synuclein appears to have similar interaction patterns with six different chaper-ones (13); outer-membrane proteins (OmpA, OmpX, FhuA) have similar properties – essentially fully unfolded – when bound to SurA and Skp chaperones (14, 15), at least when judged by their NMR fingerprint spectra. Phosphatase A displays an extended dynamic conformation, but well-defined binding poses of its interacting parts, when bound to trigger factor (16), Hsp40 (17) or SecB (18). Thus, while these reports suggest that a given protein adopts similar properties on different chaperones, the scarcity of data and absence of a direct comparison of complex structures, leaves open which interactions may confer specificity.

A pair of ‘holdase’ chaperone complexes of the mitochondrial inter-membrane space (IMS), TIM8·13 and TIM9·10, are structurally highly similar, but have different substrate binding preferences. These chaperones transport precursors of membrane proteins (henceforth denoted as ‘precursors’) to the membrane-insertase machineries in the inner membrane (TIM22) and outer mitochondrial membranes (SAM) (19)). The TIM chaperones form hetero-hexameric structures of ca. 70 kDa, composed of an alternating arrangement of Tim9 and Tim10 or Tim8 and Tim13. TIM9·10 is essential to cellular viability (20–22); even single point mutations in Tim9 or Tim10 that keep the chaperone structure intact but affect precursor protein binding can impair yeast growth and cause lethality (23). Although TIM8·13 is not essential in yeast (24), yeast cells depleted of Tim8 and Tim13 show conditional lethality (25). Additionally, mutations in the human Tim8a protein have been identified as the cause of a neurodegenerative disorder known as Mohr-Tranebjærg (MTS) or Deafness-Dystonia-Optic Neuropathy (DDON) syndrom (26, 27).

*In vivo* experiments, predominantly in yeast, have identified mitochondrial membrane proteins whose biogenesis depends on small TIM chaperones. TIM9·10 is believed to interact with all members of the mitochondrial carrier (SLC25) family, which comprises more than 50 members in humans, such as the ADP/ATP carrier (Aac in yeast); TIM9·10 furthermore transports the central components of the TIM22 and TIM23 insertion machineries (Tim23, Tim17, Tim22) as well as outer-membrane β-barrel proteins (28). TIM8·13 has a narrower clientome, and was shown to bind the precursors of the inner-membrane proteins Tim23 (25, 29, 30) and Ca^2+^-binding aspartate-glutamate carriers (31), as well as the outer-membrane β-barrel proteins VDAC/Porin, Tom40 (32) and Tob55/Sam50 (33). There is evidence that TIM8·13 does not bind the inner-membrane proteins ADP/ATP carrier (Aac) and Tim17 (25). The inner-membrane proteins that have been reported to interact with TIM8·13 have a hydrophilic domain in addition to trans-membrane domains (fig. S1), but this does not hold true for the outer-membrane β-barrels; and the potential mechanisms by which TIM8·13 may bind its clients remain unclear.

Recently, we obtained the first structure of a complex of a small TIM chaperone, TIM9·10, with the mitochondrial GDP/GTP carrier (Ggc1) (23). The structure, composed of two chaperone complexes holding one precursor protein, revealed a highly dynamic ensemble of Ggc1 conformers that form multiple short-lived and rapidly inter-converting (< 1 ms) interactions with a hydrophobic binding cleft of the chaperone (fig. S2). The TIM9·10-Ggc1 complex can be described as a “fuzzy complex", in which the high overall affinity is driven by a multitude of individually weak interactions with the hydrophobic trans-membrane (TM) parts of its clients.

To understand what confers specificity in the mitochondrial IMS chaperone system, we studied chaperone complexes of TIM9·10 and TIM8·13 with two precursor proteins, the GDP/GTP carrier (Ggc1) and the insertase component Tim23. In their native state, Ggc1 comprises six TM helices without soluble domains, and Tim23 four TM helices and a ca. 100-residue-long soluble inter-membrane space domain (Fig. 1A). By solving the complex structures of the two chaperone complexes holding Tim23, we reveal that the differential specificity of the two chaperones is based on an interplay of hydrophobic and hydrophilic interactions, which leads to different conformational properties of the precursor protein bound to these chaperones.

**Fig. 1.**
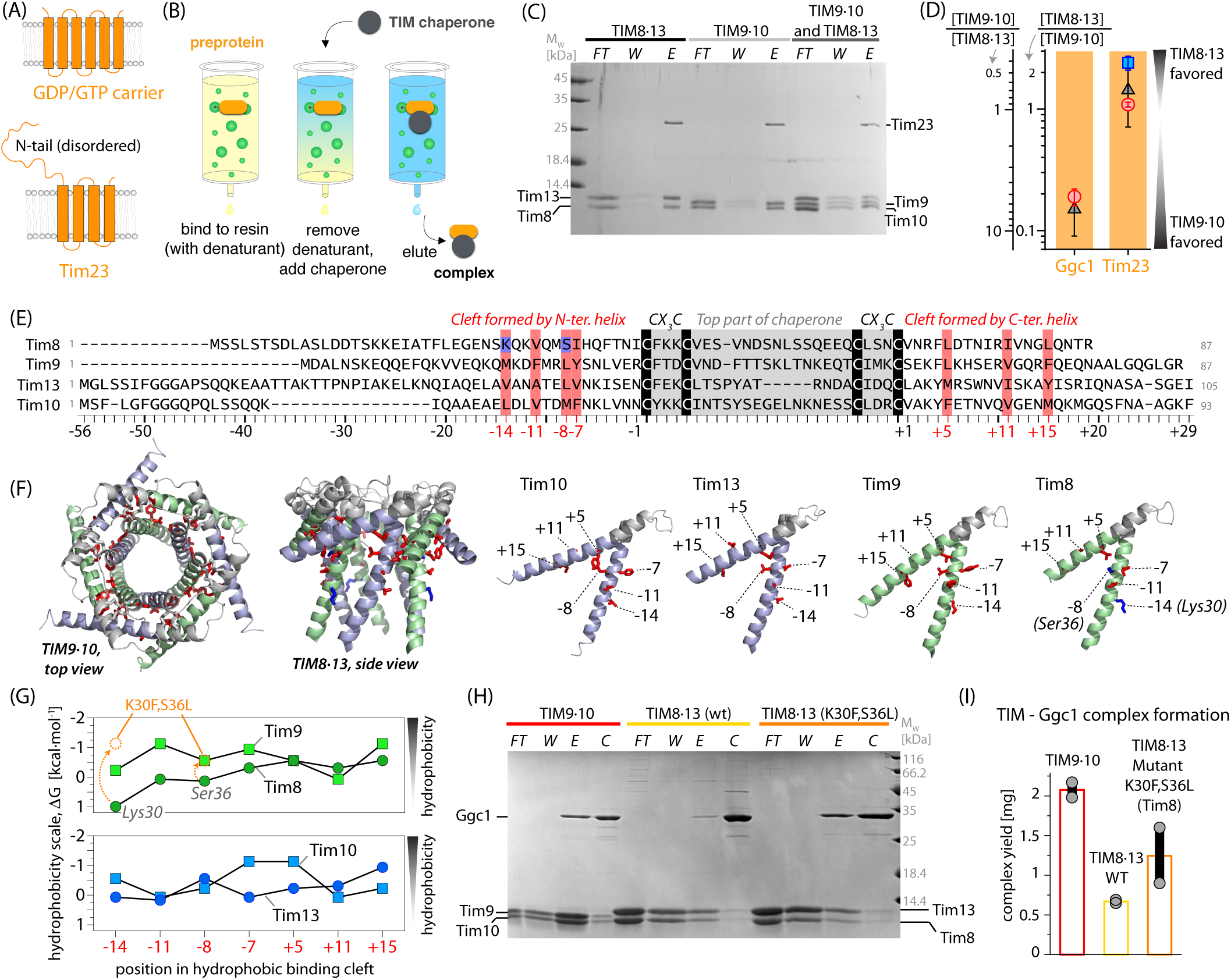
Biochemical characterization of TIM chaperone - membrane protein complexes. **(A)** Native topology of the two precursor proteins used in this study. **(B)** Schematic view of the pull-down experiment used to prepare chaperone-precursor complexes. **(C)** Formation of complexes of the two chaperones with Tim23, monitored by SDS-PAGE. In the three experiments, either TIM8·13, TIM9·10 or a 1:1 mixture of those was applied in the pull-down experiment. The lanes correspond to flow-through after applying chaperone (FT), additional wash (W), and imidazol elution (E). Because of very similar molecular weight, protein bands corresponding to Tim10 and Tim8 overlap. **(D)** Relative amounts of chaperone complexes with Ggc1 and Tim23, obtained from three different experiments: (i) a pull-down assay where both chaperones were applied to bound precursor protein (black; error bar from 3 replicates), (ii) preparation of a TIM9·10-precursor protein complex and addition of TIM8·13, and SDS-PAGE as well as mass spectrometry analysis after 1 and 3 hours (red), (iii) preparation of TIM8·13-Tim23 followed by TIM9·10 addition and SDS-PAGE as in (ii). Error estimates were obtained from two experiments. The protein amounts were determined from LC/ESI-TOF mass spectrometry (fig. S4). On the two axis are the ratios of TIM9·10 over TIM8·13 (left) or its inverse (right). **(E)** Sequence alignment of the four small Tims, using a numbering starting at the conserved Cys residues towards the N-and C-termini. Highlighted in blue are Tim8 residues mutated to have more hydrophobic TIM8·13(Tim8_K30F,S36L_). Conserved hydrophobic residues are highlighted in red throughout this manuscript. **(F)** Location of the residues in the hydrophobic cleft. **(G)** Comparison of Kyte-Doolittle hydrophobicity of the residues in the binding cleft of wild-type (WT) native Tim proteins and Tim8_K30F,S36L_. **(H)** Pull-down experiment of Ggc1 with TIM9·10, TIM8·13 and TIM8·13(Tim8_K30F,S36L_). Lane descriptions are as in (C); additionally, the fraction obtained after final wash with guanidine hydrochloride and imidazol, to control the Ggc1 initially loaded onto the column, is shown (control, C). **(I)** Amount of complex obtained from pull-down experiments of WT and mutant chaperones; the same amount of Ggc1 was applied in all three experiments, and the total amount of eluted complex was determined spectroscopically.

## Results

### TIM8·13 and TIM9·10 interact differently with membrane precursor proteins

We have developed an experimental protocol (23) to prepare complexes of the inherently insoluble membrane-protein precursors and chaperones (Fig. 1B,C). Briefly, the approach involves the recombinant production of His-tagged precursor protein, binding it to a His-affinity column in denaturing conditions, followed by removal of the denaturant and simultaneous addition of a chaperone. The chaperone-precursor complex is then eluted for further biochemical, biophysical and structural investigations.

The measurement of dissociation constants of chaperones and membrane precursor proteins, using methods such as isothermal titration calorimetry or surface plasmon resonance, is not possible, because the complexes cannot be formed in solution (e.g. flash-dilution methods, which work for other chaperones (14), failed; data not shown). Thus, to characterize the relative affinities of the precursor proteins to the two chaperones, we performed different types of competition experiments. In a first experiment, precursor protein was bound to the resin, and both chaperones were simultaneously added, before washing excess chaperone, and eluting the chaperone-precursor complexes (Fig. 1C). NMR spectroscopy shows that the two chaperones do not form mixed hetero-hexameric complexes, implying that TIM9·10 and TIM8·13 stay intact in such competition experiments (fig. S3). In a second class of experiments, we prepared one type of complex (e.g. TIM9·10-Tim23) and added the other chaperone (e.g. TIM8·13) in its apo state, allowing the precursor protein to be transferred. These experiments also demonstrate that membrane precursor proteins can be transferred between these two chaperones, on the time scale we investigated (minutes to hours). We used SDS-PAGE analyses and electrospray ionization mass spectrometry (ESI-MS) to systematically quantify the amount of obtained complexes (Fig. 1D and fig. S4). Consistently, we find that Ggc1 has a strong preference for TIM9·10 (ca. 5-to 10-fold), while Tim23 shows a slight preference for TIM8·13 (ca. 1.5-fold). Taken together, we established that the two chaperones bind with different affinities to two inner-membrane precursor proteins, whereby TIM8·13 is barely able to hold Ggc1, in contrast to TIM9·10, while it can hold Tim23 slightly better than TIM9·10.

### The small TIM chaperones use a conserved hydrophobic cleft for membrane precursor protein binding

To understand the different binding properties, we performed a sequence alignment of the small TIMs, which reveals a well conserved set of hydrophobic residues that point towards the binding cleft formed between the inner (N-terminal) and outer tentacles (23) (Fig. 1E,F). The overall hydrophobicity of these residues is lower in Tim8 and Tim13 than in Tim9 and Tim10 (Fig. 1G). In particular, Tim8 has a charged residue in position −14 (K30) and a polar one in position −8 (S36) of the hydrophobic motif. (The sequences are numbered starting with negative numbering at the twin CX_3_C motif towards the N-terminus, and positive numbering from the last Cys to the C terminus). We speculated that the less hydrophobic nature of TIM8·13’s binding cleft reduces its affinity to TM parts of membrane precursor proteins. To test this hypothesis, we generated a mutant TIM8·13 with increased hydrophobicity (Tim8_K30F,S36L_; Fig. 1G). This more hydrophobic TIM8·13(Tim8_K30F,S36L_) complex allows obtaining significantly larger amounts of complex with Ggc1 than native TIM8·13 under otherwise identical conditions (Fig. 1H,I). This observation establishes the importance of the hydrophobic cleft for binding hydrophobic TM parts of precursor proteins.

To better understand the client-binding properties of the two chaperones, we turned to structural studies. Solution-NMR spectra of apo TIM8·13 (Fig. 2A) and residue-wise resonance assignments allowed identifying the residues forming secondary structure and estimating their local flexibility. In agreement with the crystal structure, the core of rather rigid tentacles comprises the top part of the chaperone between the CX_3_C motifs and ca. 15-25 residues before and after these motifs, while about 10-20 residues on the N-and C-termini are flexible (fig. S5). To probe the binding of a trans-membrane segment of a membrane precursor protein, we performed NMR-detected titration experiments of TIM8·13 with a cyclic peptide corresponding to the two C-terminal strands of the β-barrel voltage-dependent anion channel (VDAC_257-279_) that has a propensity to form a β-turn (35). Addition of this cyclic VDAC_257-279_ induces chemical-shift perturbations (Fig. 2B), that are primarily located in the hydrophobic cleft formed between the inner and the outer rings of helices (Fig. 2C,D). This binding site matches very closely the site on TIM9·10 to which VDAC_257-279_ binds (23) (Fig. 2C). Interestingly, the VDAC_257-279_-induced chemical-shift perturbation (CSP) effects in TIM8·13 are overall only about half of the magnitude of CSPs found in TIM9·10, pointing to a higher population of TIM9·10-VDAC_257-279_ complex compared to TIM8·13-VDAC_257-279_ at comparable conditions (Fig. 2C). This finding suggests a lower affinity of TIM8·13 to VDAC_257-279_, as expected from its lower hydrophobicity of TIM8·13, compared to TIM9·10. Photo-induced cross-linking experiments of a Bpa-modified VDAC_257-279_ peptide to TIM8·13 show that only the cyclic peptide forms cross-linking adducts while the linear, mostly disordered (35) form does not (Fig. 2F), mirroring identical findings with TIM9·10 (23) and yeast cytosolic chaperones Ssa1, Ydj1, Djp1, and Hsp104 (36). A rationale for this finding is the fact that in a β-turn the side chains of consecutive residues point to the two opposing faces thus creating one hydrophobic and one more hydrophilic face (Fig. 2E). In contrast, due to its disorder, the linear VDAC_257-279_ peptide does not have a stable hydrophobic face, reducing its affinity to the hydrophobic binding cleft on the chaperone.

**Fig. 2.**
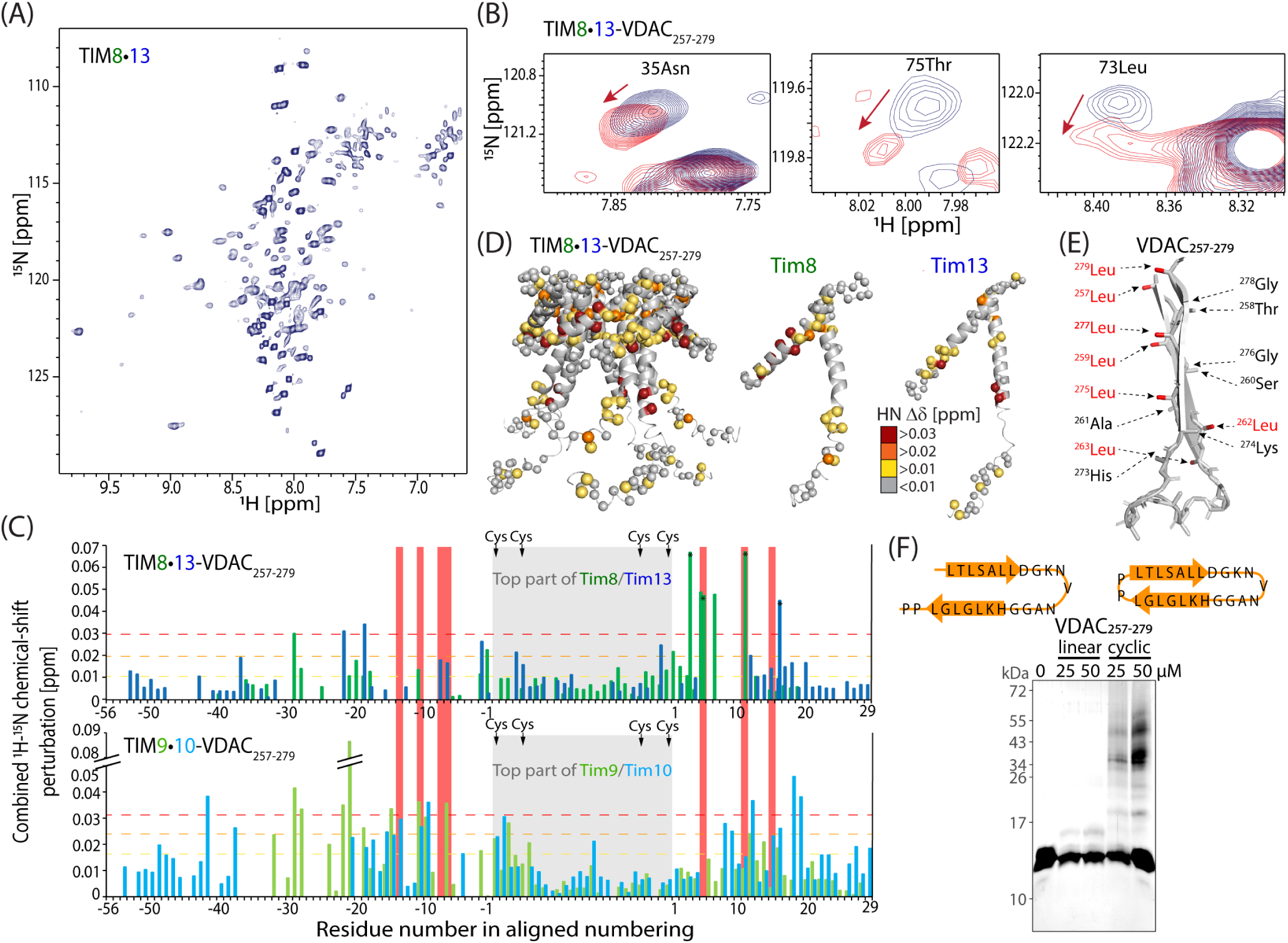
Solution-NMR and binding of a VDAC fragment to TIM8·13. **(A)** ^1^H-^15^N NMR spectrum of TIM8·13 at 35 °C. **(B)** Chemical-shift perturbation (CSP) in TIM8·13 upon addition of 5 molar equivalents of VDAC_257-279_. **(C)** CSP effects of VDAC_257-279_ binding. The data for TIM9·10 are from ref. (23). **(D)** Plot of CSP data on the TIM8·13 structure. **(E)** Schematic structure of the two last strands of VDAC, as found in the NMR structure (34) of the full β-barrel, showing that the hydrophobic and hydrophilic side chains cluster on the two opposite faces of the β-turn. **(F)** Photo-induced cross-linking of the linear (left) and cyclic (right) VDAC_257-279_ peptides.

Collectively, these data establish that both chaperones use the same conserved binding cleft to interact with hydrophobic membrane precursor protein sequences, and that TIM9·10, presumably due to the higher hydrophobicity of its binding cleft, interacts more efficiently with TM parts, and thus with Ggc1 and the VDAC fragment. In light of this observation, how does TIM8·13 achieve a binding affinity to Tim23 which is slightly higher than the one of TIM9·10 (Fig. 1D)?

### Hydrophilic fragments interact differently with TIM8·13 and TIM9·10

Tim23 has a hydrophilic N-terminal segment in addition to four TM helices (Fig. 3A), and we investigated whether this part interacts with the chaperones. NMR spectra of the soluble Tim23_IMS_ fragment (residues 1-98) in isolation show the hallmark features of a highly flexible intrinsically disordered protein with low spectral dispersion of ^1^H-^15^N NMR signals (Fig. 3B,C, orange spectrum), as previously reported (37). Upon addition of TIM9·10, the Tim23_IMS_ ^1^H-^15^N spectrum (Fig. 3B, left) shows only small changes: all cross-peaks are still detectable, and small chemical-shift perturbations (CSPs) are only observed for a few residues at the N-terminus, which has higher hydrophobicity (Fig. 3D). This finding suggests only very weak, possibly non-specific interactions between the very N-terminus of Tim23_IMS_ and TIM9·10. In line with this finding, the interaction is not detectable by isothermal titration calorimetry (ITC) measurements (Fig. 3H).

**Fig. 3.**
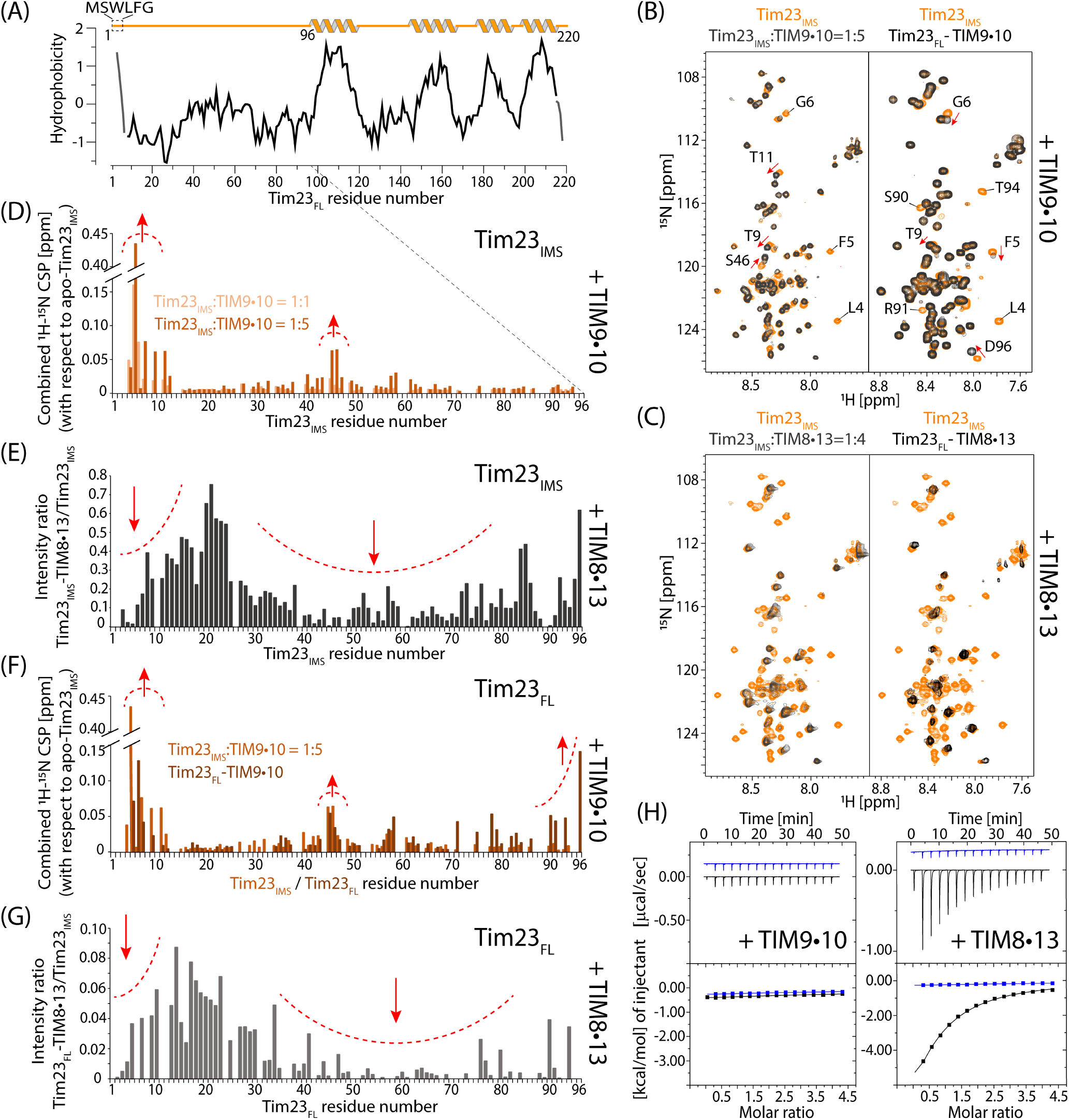
Tim23 has markedly different properties when binding to TIM8·13 and to TIM9·10. **(A)** Hydrophobicity of Tim23 (Kyte-Doolittle). **(B)** NMR spectra of the ^15^N-labeled soluble Tim23_IMS_ fragment in the presence of TIM9·10 (left, black), and of full-length Tim23 bound to TIM9·10 (right, black) are compared to the Tim23_IMS_ fragment in isolation (orange), under identical buffer conditions and NMR parameters. **(C)** As in (B) but with TIM8·13 instead of TIM9·10. **(D)** Chemical-shift perturbation (CSP) of residues in Tim23_IMS_ upon addition of 1 (light orange) or 5 (dark orange) molar equivalents of TIM9·10. **(E)** Intensity ratio of residues in Tim23_IMS_ in the presence of 4 molar equivalents of TIM8·13 compared to Tim23_IMS_ alone. **(F)** CSP of the detectable residues in full-length Tim23 attached to TIM9·10 (brown), compared to the soluble Tim23_IMS_ fragment. **(G)** Intensity ratio of detectable residues in Tim23_FL_ attached to TIM8·13. Note that the ratio was not corrected for differences in sample concentration, and the scale cannot be compared to the one in panel (E). **(H)** Calorimetric titrations for the interaction of TIM9·10 or TIM8·13 (54 µM in the calorimetric cell) with Tim23_IMS_ (1.15 mM in the injecting syringe). Thermograms (thermal power as a function of time) are displayed in the upper plots, and binding isotherms (ligand-normalized heat effects per injection as a function of the molar ratio, [Tim23_IMS_]/[chaperone]) are displayed in the lower plots. Control experiments, injecting into a buffer, are shown in blue.

The interaction of the hydrophilic Tim23_IMS_ fragment with TIM8·13 is significantly stronger, with pronounced binding effects detected by ITC, and a dissociation constant of K_*d*_ = 66 ±8 µM (Fig. 3H, right panel). The ^1^H-^15^N NMR spectrum of Tim23_IMS_ in the presence of TIM8·13 shows strongly reduced peak intensities for the majority of the residues (Fig. 3C, left). Such a peak broadening is expected when a highly flexible polypeptide binds to a relatively large object such as TIM8·13, thereby inducing faster nuclear spin relaxation and thus broader signals of lower magnitude. Analysis of the peak-intensity reduction reveals two regions of Tim23 that are particularly involved in the binding: (i) the N-terminal hydrophobic residues, which are also involved in interacting with TIM9·10, and (ii) a long sequence stretch comprising residues from ca. 30 to 80 (Fig. 3E).

To characterize the conformation of full-length (FL) Tim23 bound to TIM8·13 and TIM9·10, we prepared Tim23_FL_-labeled Tim23–chaperone complexes using the method outlined in Fig. 1A. Very similarly to the experiments with the Tim23_IMS_ fragment, the signals corresponding to the N-terminal half of Tim23 are still intensely visible in the Tim23-TIM9·10 complex. The small observed CSPs are localized primarily at the ten N-terminal residues (Fig. 3B,F). In contrast, when Tim23_FL_ is bound to TIM8·13, the signals corresponding to its N-terminal half are severely reduced in intensity, revealing tight contact of the flexible N-terminal half of Tim23 to TIM8·13 (Fig. 3C,G).

In neither of the two Tim23_FL_ complexes any additional signals, that may correspond to Tim23’s TM helices, are visible. We ascribe this lack of detectable signals of residues in the TM part to extensive line broadening. The origin of this line broadening may be ascribed to the large size of the complex and likely to additional millisecond (ms) time scale dynamics of Tim23’s TM parts in the hydrophobic binding cleft of the chaperones. Such millisecond motions have been found in the TIM9·10–Ggc1 complex (23).

### TIM8·13 uses an additional hydrophilic face for protein binding

We probed the binding sites that the chaperones use to interact with Tim23_IMS_ or Tim23_FL_ using NMR spectroscopy on samples in which only the chaperone was isotope-labeled. Interestingly, the CSPs in the two chaperones upon addition of Tim23_IMS_ reveal distinct binding patterns (Fig. 4A): in TIM8·13, the largest effects involve residues in the hydrophilic top part of the chaperone, between the CX_3_C motifs, as well as a few residues toward the C-terminal outer ring of helices; in contrast, the corresponding top part of TIM9·10 does not show any significant effects, but CSPs are observed at residues in the hydrophobic binding cleft, and in particular the N-terminal helix (Fig. 4B). This data, together with the Tim23_IMS_-detected data in Fig. 3 establish that TIM8·13 uses its hydrophilic top part to bind Tim23’s N-terminal half, while only a short stretch of hydrophobic residues at the very N-terminus of Tim23 interacts with the hydrophobic cleft of TIM9·10, which is also the binding site of TM parts (Figs. 1 and 2).

**Fig. 4.**
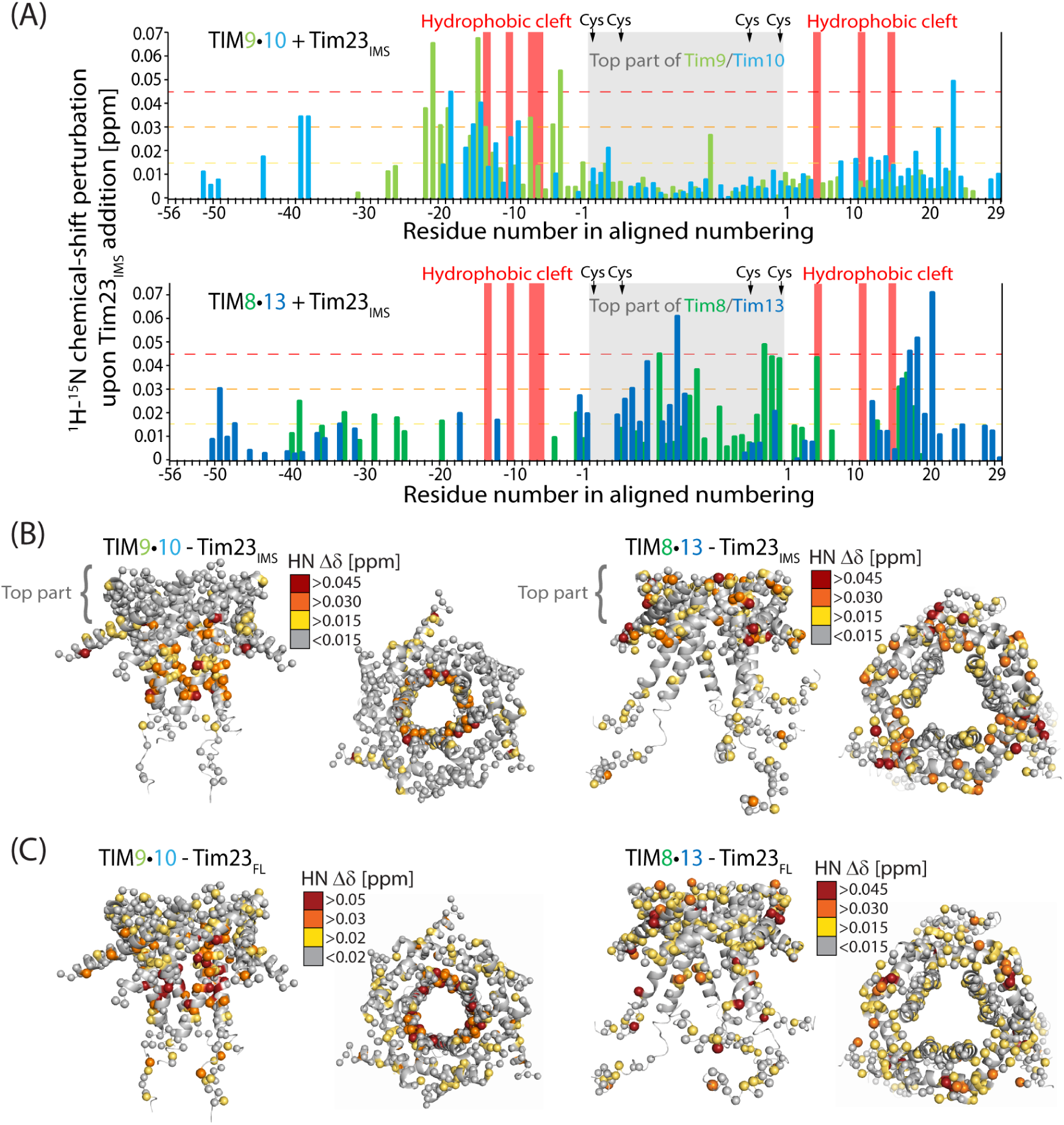
Tim23_IMS_ and full-length Tim23 differ in their interactions with TIM9·10 and TIM8·13 chaperones. **(A)** Chemical-shift perturbations observed upon addition of the Tim23_IMS_ fragment to TIM9·10 (top) and TIM8·13 (bottom). The chaperone:Tim23_IMS_ ratios were 1:1 (TIM8·13) and 1:3 (TIM9·10). **(B)** Mapping of Tim23_IMS_-induced CSPs on TIM9·10 and TIM8·13, showing that while the top part of TIM9·10 does not show any significant CSPs, the corresponding part is the main interacting region of TIM8·13. **(C)** CSP in complexes of TIM9·10 (left) and TIM8·13 (right) bound to full-length Tim23.

Chaperone-labeled complexes with Tim23_FL_ confirm these findings, and point to the additional effects induced by the bound TM part: in TIM9·10-Tim23_FL_, large CSP effects are located primarily in the binding cleft, in line with the view that the top part of TIM9·10 is not involved in binding Tim23. In contrast, Tim23_FL_-induced CSPs are found across the whole TIM8·13, including the hydrophilic top and the hydrophobic cleft (Fig. 4C and fig. S6).

Collectively, NMR, ITC and mutagenesis have revealed that the hydrophobic cleft of both TIM8·13 and TIM9·10 are essential to hold the hydrophobic parts of the clients, and that TIM8·13, but not TIM9·10, additionally interacts with the hydrophilic part of Tim23 to increase its affinity. This interaction, which is mediated by the hydrophilic top part of TIM8·13, reduces the conformational flexibility of Tim23’s N-terminal half. This hydrophilically driven interaction supports previous observation during the protein import (25), where TIM8·13 interacting with hydrophobic membrane precursor without long hydrophilic stretches (fig. S1) was detected only when they were fused to hydrophilic Tim23_IMS_.

### Structural ensembles of chaperone-Tim23 complexes

We integrated the NMR data with further biophysical, structural and numerical techniques to obtain a full structural and dynamical description of the complexes. We first investigated the complex stoichiometry using multi-angle light scattering (SEC-MALS), NMR-detected diffusion-coefficient measurements, and analytical ultra-centrifugation. These methods, which provide estimates of molecular mass (and shape) from orthogonal physical properties (gel filtration and light scattering; translational diffusion), reveal properties best compatible with a 1:1 (chaperone:precursor) stoichiometry (Fig. 5A and fig. S7). This stoichiometry contrasts the 2:1 (chaperone:precursor) stoichiometry for TIM9·10 holding the 35 kDa-large carrier Ggc1 (23) (fig. S2). Small-angle X-ray scattering (SAXS) data of both TIM9·10-Tim23 and TIM8·13-Tim23 also point to a molecular weight corresponding to a 1:1 complex (SAXS; Fig. 5B). Importantly, SAXS provides significantly more information, namely the overall shape of the ensemble of conformations present in solution. As our data point to large-scale flexibility, in particular of the membrane precursor protein, this data is best analyzed by considering the dynamic ensemble explicitly. We used molecular dynamics simulations to account for the breadth of possible conformations that, collectively, give rise to the observed scattering. To effectively sample the conformational space of the chaperone-Tim23 complex, we constructed two distinct structural models in which the N-terminal half of Tim23 is either modeled as a floppy unstructured tail or bound to the hydrophilic upper part of chaperone, corresponding to the ‘N-tail unbound’ and ‘N-tail bound’ conformation, respectively. In both models, the hydrophobic Cterminal transmembrane domain of Tim23 is bound to the hydrophobic cleft of the chaperone, as identified by NMR (Fig. 4 and fig. S6C,D). Initiating from both conformations, explicit solvent atomistic MD simulations (∼four microseconds in total) were performed to collect the structures for the ‘N-tail unbound’ and ‘N-tail bound’ ensembles. In the case of TIM8·13, the ensemble of ‘N-tail bound’ state recapitulates the experimentally observed pattern better than the ensemble of ‘N-tail unbound’ state (Fig. 3 C and E). We then constructed a mixed ensemble consisting of both ‘Ntail bound’, and ‘N-tail unbound’ states which were used for further ensemble refinement using the Bayasian Maximum Entropy (BME) method guided by experimental SAXS data (23, 38, 39). We found that the experimental SAXS data of TIM8·13–Tim23 are very well reproduced when the mixed ensemble has >85 % of the ‘N-tail bound’ state (Fig. 5c, D). In contrast, the experimental data of TIM9·10–Tim23 are only well reproduced when the TIM9·10–Tim23 ensemble comprises >75 % of the ‘N-tail unbound’ state. This refined ensembles guided by experimental SAXS data are in excellent agreement with the NMR data, which showed that (i) in the TIM9·10-Tim23 complex, the N-terminal part of Tim23 is predominantly free and flexible, and Tim23 makes contacts only to the hydrophobic cleft of the chaperone, while (ii) in TIM8·13-Tim23, the Tim23_Nter_ part is largely bound to the upper part of the chaperone (Figs. 3 and 4).

**Fig. 5.**
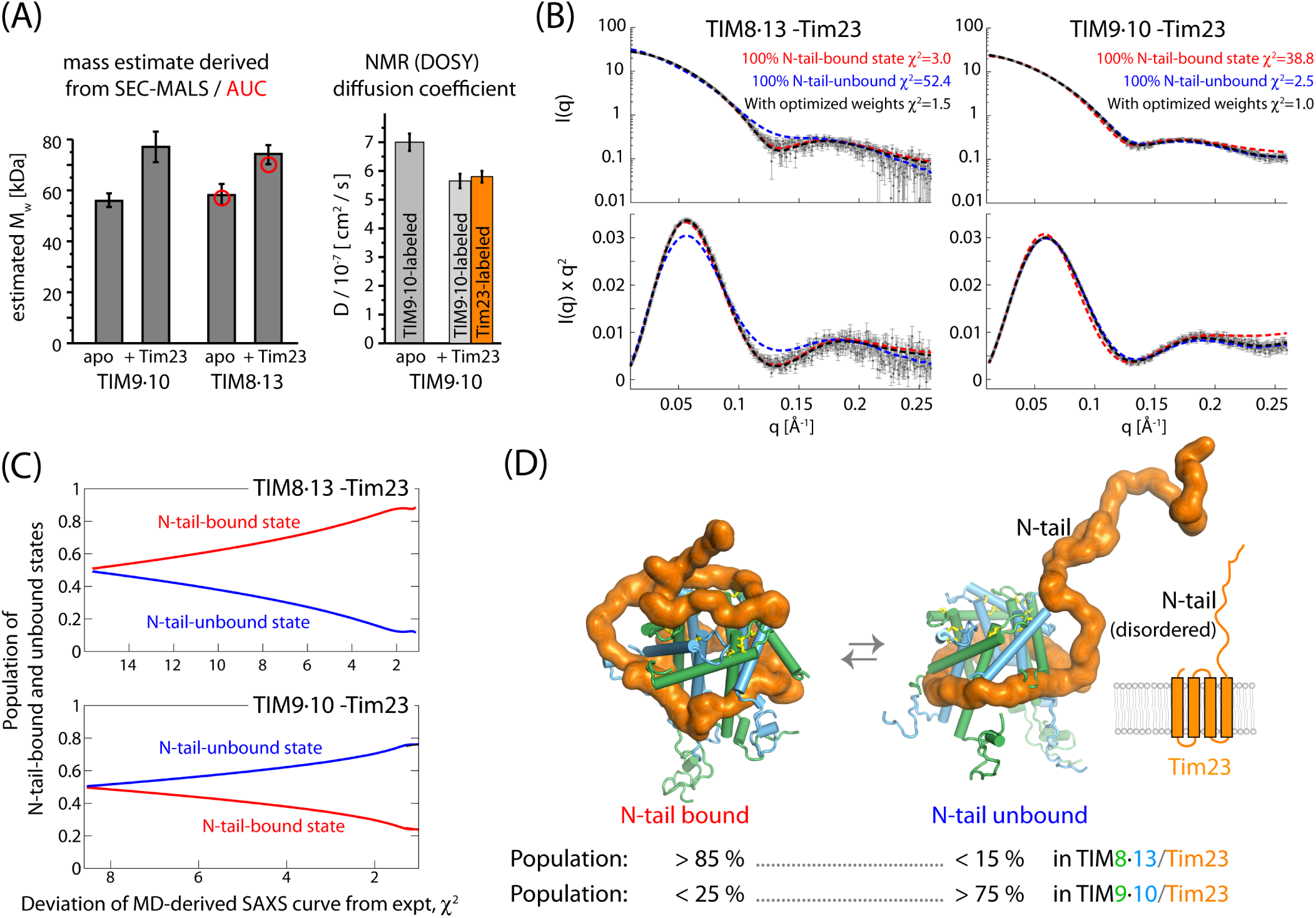
Architecture of the TIM8·13 and TIM9·10 holdases in complex with full-length Tim23. **(A)** (Left) Apparent molecular weights of apo and holo chaperone complexes from SEC-MALS, and AUC (red) circles. (Right) Translational diffusion coefficients of TIM9·10 (apo) and TIM9·10-Tim23_FL_ from NMR DOSY measurements. Two independent samples were used for the complex, in which either the chaperone or the precursor protein was labeled, as indicated. See also fig. S7. **(B)** Small-angle X-ray scattering curves (top) and Kratky plot representations thereof for the two chaperone-precursor complexes. The lines are SAXS curves calculated from structural ensembles obtained over xx µs long MD trajectories, in which the N-terminal half of Tim23 was either in a conformation bound to the top part of the chaperone (red) or in a loose unbound conformation (blue), or from an ensemble in which these two classes of states were present with optimized weights. **(C)** Goodness of fit of the back-calculated SAXS curves to the experimental SAXS data as a function of the relative weights of the two classes of conformations, in which the N-terminal half of Tim23 is either bound or unbound. **(D)** Snapshots of conformations in which Tim23_N-tail_ is either bound or unbound, and the best-fit relative weights of the two classes of states as derived from SAXS/MD. More SAXS/MD data and ensemble views are provided in fig. S8.

The amount of ‘N-tail bound’ relative to ‘N-tail unbound’ states is expected to depend on the affinity of the N-terminus of Tim23 to the chaperone. Indeed, the ITC-derived TIM8·13-Tim23_IMS_ affinity (K_d_=66 µM; Fig. 3H) predicts that the population of N-tail-bound states is of the order of 75 - 98 % (see Methods for details), in excellent agreement with the MD/SAXS derived value (> 85 %). This good match of data from the Tim23_IMS_ fragment and Tim23_FL_ suggests that binding of Tim23’s hydrophilic domain is similar in the fulllength complex. Similarly, the inability to detect TIM9·10-Tim23_IMS_ binding by ITC is mirrored by the small population of the ‘N-tail bound’ state in the full-length complex.

To identify the molecular mechanisms underlying the observed differences in N-tail binding, we studied the interactions formed between Tim23 and the chaperones. An interesting pattern emerges from analysis of the electrostatic interactions (Fig. 6A). Tim23 contributes primarily with positively charged residues, which contact negatively charged residues on the chaperones. For example, three key aspartate or glutamate residues are involved in binding of lysine or arginine residues of Tim23_IMS_. The top part of TIM8·13 has predominantly polar and negatively charged residues, which are in transient contact with the positive charges of Tim23 N-tail, within a dynamic ensemble of conformations (Fig. 6B,C). Interestingly, in Tim9, a lysine (K51) contributes a positive charge, where the equivalent position in TIM8·13 is a non-charged, polar residue. We propose that the less complementary electrostatic properties of TIM9·10’s top part and Tim23’s N-tail, as compared to TIM8·13, diminish the affinity of the N-tail to TIM9·10, thus explaining the low probability of finding the N-tail-bound state in the TIM9·10 complex.

**Fig. 6.**
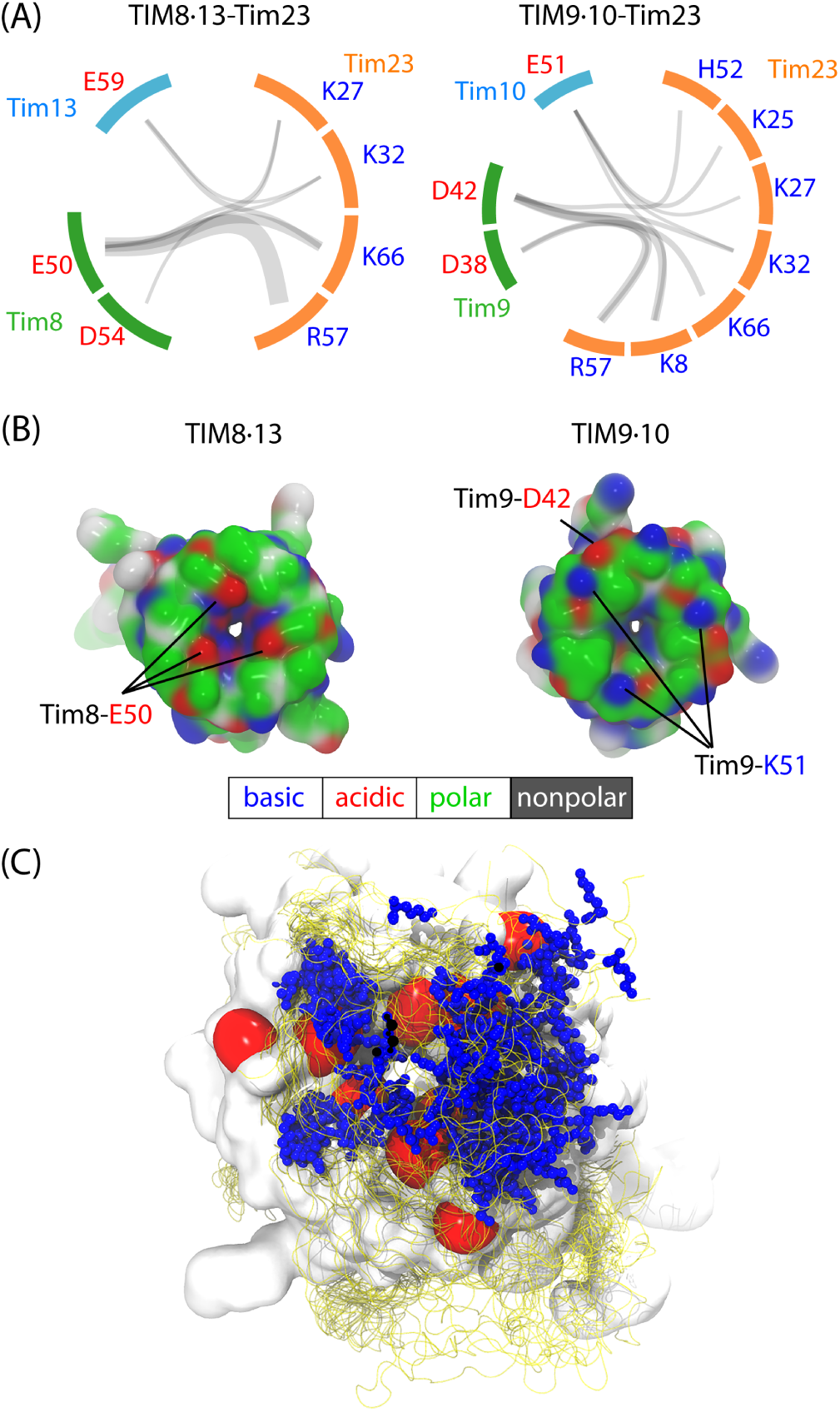
Charged interactions that drive the N-tail-chaperone interaction. **(A)** Principal electrostatic interactions between the N-tail of Tim23 and the chaperones within the ensemble of ‘N-tail bound’ conformations. The charged residue pairs forming salt bridges are connected by grey semi-transparent lines whose thickness linearly scales with the frequency of the corresponding salt bridge observed in MD simulations. Despite more diverse salt bridges were observed in TIM9·10–Tim23 (10 in TIM9·10–Tim23 and 7 in TIM8·13–Tim23), these salt bridges were in average less stable than the ones in TIM8·13–Tim23, likely resulting in overall weaker interactions. **(B)** Snapshots of top views of the two chaperones along MD simulations of their holo forms in complex with TIM23. The top views of the chaperones in the apo forms are shown in Fig. S5E and F. Residues are color-coded according to the scheme reflected below the figure. **(C)** Ensemble view of the N-tail-bound state of TIM8·13–Tim23. Red surface represents the negatively charged E59 of Tim13 and E50 and D54 of Tim8. Blue stick-and-ball represents the sidechain of positively charged residues (K8, K25, K27, K32, R57 and K66) of Tim23, which is shown as an ensemble of 25 structures. Ensemble view of the N-tail-bound state of TIM9·10–Tim23 is shown in Fig. S5G.

## Discussion

Transfer chaperones (holdases) need to fulfill two contradicting requirements, holding their clients very tightly to avoid their premature release and aggregation, while at the same time allowing release at the downstream factor. This apparent contradiction is solved by a subtle balance of multiple individually weak interactions, and a resulting dynamic complex, wherein the precursor protein extensively samples a wide range of different conformations. This ensemble of conformations results in a high overall affinity, yet a downstream foldase/insertase can detach the precursor protein from the chaperone without significant energy barrier. Balancing the interaction strength is, thus, crucial to chaperone function, and highlighted by the fact that single-point mutations in TIM9·10 can abrogate the client binding (23). Herein, we have revealed a fine-tuning of chaperone-client specificity that involves hydrophobic interaction with the chaperone’s binding cleft and additional hydrophilic interactions, mostly mediated by charged residues, with the chaperone’s top part. Lower hydrophobicity within the binding cleft of TIM8·13 compared to TIM9·10 arises by overall less hydrophobic residues and a positively charged residue (Lys/Arg) that is highly conserved in Tim8. As a consequence, TIM8·13 is less able to hold the TM parts of its clients than TIM9·10 by ca. one order of magnitude. As we showed, replacement of two charged/polar side chains in TIM8·13’s cleft brings TIM8·13 almost to the same level as TIM9·10 for holding an all-transmembrane client.

For binding of its native client Tim23, TIM8·13 uses additional hydrophilic interactions to its client’s IMS segment, which, as we show here, is ineffective in TIM9·10-Tim23 interaction. The additional interaction effectively compensates for the lower affinity of TIM8·13 to the client’s TM part. In the case of Tim23, this additional interaction involves a sequence stretch of at least 35-40 residues. Interestingly, TIM8·13 has also been shown to be involved in the transport of a Ca^2+^-regulated mitochondrial carrier, the Asp/Glu carrier with an extended hydrophilic domain (fig. S1). Interestingly, membrane precursor proteins that have been shown *not* to interact with TIM8·13, such as mitochondrial carriers (Ggc, Aac) and Tim17, do not have hydrophilic stretches longer than ca. 10-15 residues (fig. S1), suggesting a minimum length larger than about 20-25 residues required for binding.

The nature of these additional hydrophilic interactions appear to involve primarily positively charged residues on the precursor protein that contact negative (Asp/Glu) residues located on the top part of the chaperone. The presence of a positively charged residue, K51 of Tim9, is presumably contributing to the low probability of finding Tim23’s N-tail bound to TIM9·10.

This study provides a rationale why mitochondria contain two very similar IMS chaperone complexes, the essential TIM9·10 and the non-essential TIM8·13 complex. The current results propose that for some substrates (like Tim23, or Asp-Glu carrier; see Figure S1), TIM8·13 can contribute stabilizing interactions with the hydrophilic soluble parts. The observation that this dual system is conserved even in human suggests that the presence of the TIM8·13 system is not just the result of gene duplication, which appear rather often in yeast. Our study has also revealed that mitochondrial membrane precursor proteins may be transferred from one chaperone to the other, opening the possibility that these two chaperones truly cooperate in precursor protein transfer to downstream insertases.

Taken together, our study reveals how a subtle balance of hydrophobic and hydrophilic interactions is used to tune promiscuity versus specificity in molecular chaperones. We propose that a similar balance of interactions determines the clientome of the cellular chaperones.

## Methods

### Plasmids

Genes coding for *Saccharomyces cerevisiae* Tim8 and Tim13 were cloned in the co-expression plasmid pETDuet1. The expressed protein sequences were MSSLST SDLASLDDTSKKEIATFLEGENSKQKVQMSIHQFTNI CFKKCVESVNDSNLSSQEEQCLSNCVNRFLDTNIRIV NGLQNTR (Tim8) and MGSSHHHHHHSQDPSQDPEN LYFQGGLSSIFGGGAPSQQKEAATTAKTTPNPIAKEL KNQIAQELAVANATELVNKISENCFEKCLTSPYATR NDACIDQCLAKYMRSWNVISKAYISRIQNASASGEI (Tim13). A tobacco etch virus (TEV) cleavage site on Tim13 allows generating a final construct starting with GGLSS (the native Tim13 sequence starts with MGLSS). The same approach was used for preparing TIM9·10, including coexpression of the two proteins, with a cleavable His_6_ tag on one of the proteins (Tim10), as described elsewhere (23). The gene coding for full-length *S. cerevisiae* Tim23 (C98S, C209S, C213A) with a C-terminal His_6_-tag was cloned in the expression plasmid pET21b(+). The plasmid for expression of the intrinsically disordered N-terminal domain of *S. cerevisiae* Tim23_*IMS*_ (residues 1 to 98), with a Nterminal glutathion S-transferase (GST) tag, was a gift from Markus Zweckstetter’s lab and described earlier (37). The *S. cerevisiae* Ggc1(C222S) construct was designed with a Cterminal 6xHis-tag in pET21a expression plasmid, reported earlier (23).

### Protein expression and purification

We found that chaperone complexes of TIM9·10 and TIM8·13 can be obtained either by over-expression in SHuffle T7 or BL21(DE3) *Escherichia coli*. Expression in the former results in soluble protein with correctly formed disulfide bonds, while the latter requires refolding from inclusion bodies. The proteins obtained with either method have indistinguishable properties (SEC, NMR). For TIM9·10, SHuffle expression results in better yield, while we obtain higher TIM8·13 yields with refolding from BL21(DE3). Accordingly, TIM9·10 and unlabelled TIM8·13 were over-expressed in the SHuffle T7 *Escherichia coli* cells and purified as described previously (23). Over-expression of the isotope labelled TIM8·13 chaperone complex from the BL21(DE3) *E. coli* cells was induced with 1 mM IPTG, and the cells were incubated for 4 hours at 37ºC. Cell pellets were sonicated and the inclusion-body fraction was resuspended sequentially, first in buffer A (50 mM Tris((tris(hydroxymethyl)aminomethane), 150 mM NaCl, pH 7.4) supplemented with 1% LDAO and 1% Triton100, then in buffer A supplemented with 1M NaCl and 1M urea and lastly in buffer B (50 mM Tris, 250 mM NaCl, pH 8.5). The last pellet fraction was solubilized in buffer B supplemented with 50 mM DTT and 3 M guanidine-HCl at 4ºC over night. The TIM8·13 complex was refolded by rapid dilution in buffer B containing 5 mM glutathione (GSH) and 0.5 mM glutathione disulfide (GSSG). The complex was purified on a NiNTA affinity column and the affinity tag was removed with TEV protease and an additional NiNTA purification step. Full-length precursor proteins, Tim23 and Ggc1, were expressed as inclusion bodies from BL21(DE3) cells, at 37°C during 1.5 and 3 hours, respectively, after adding 1 mM IPTG. Precursor proteins were solubilized in buffer A supplemented with 4 M guanidine-HCl for Tim23 and 6 M guanidine-HCl for Ggc1, at 4ºC over the night. Precursor proteins were purified by affinity chromatography, in the same denaturating conditions used for solubilization. Imidazole was removed from the precursor protein sample with dialysis in buffer A supplemented with 4 M guanidine-HCl. GST-tagged Tim23_IMS_ was expressed in the soluble protein fraction from BL21(DE3)Ril+ cells during 4 h at 25°C, after adding 1 mM IPTG. After sonication of the cell pellets, the soluble protein fraction was incubated with glutathion-agarose resin for 2 hours at 4ºC. After washing the unspecifically bound proteins with 10 CV of Buffer A, the GST-tag was cleaved from the Tim23_IMS_ by incubating the resin with 1 mg of TEV protease per 50 mg of precursor protein, at 4ºC over the night. Cleaved Tim23_IMS_ and the protease were collected in the flow-through and an additional NiNTA purification step was applied to remove the TEV protease from the protein sample. Soluble Tim23_IMS_ was subjected to gel filtration on a Superdex 75 10/300 column and stored in buffer A.

For NMR experiments proteins were expressed in D_2_O M9 minimal medium and labeled either with 1 g/L ^15^NH_4_Cl and 2 g/L D-[^2^H,^13^C] glucose or specifically labeled on isoleucine, alanine, leucine, valine side chains using the QLAM-A^*β*^I^*δ*1^L^*proR*^V^*proR*^ kit from NMR-Bio (www.nmr-bio.com) according to the manufacturer’s instructions.

The fragments of human VDAC1 peptide (cyclic or linear VDAC_257-279_) were prepared by solid-phase synthesis as described elsewhere (35), lyophilized, and resolubilized first in DMSO and then stepwise dilution to buffer, as described elsewhere (23). The peptide used for photo-induced cross-linking differed from the one used for NMR by the substitution of L263 by a Bpa side chain, as used earlier (23, 35).

### Preparation of chaperone-precursor protein complexes

Purified full-length precursor protein, Tim23 or Ggc1, was bound to NiNTA resin in 4 M guanidine-HCl. The column was washed with five column volumes (CV) of buffer A supplemented with 4 M guanidin-HCl, and with 5 CV of buffer A. A twofold excess of the chaperone complex was passed through the column twice. The column was washed with 10 CV of buffer A and the precursor-chaperone complex was eluted in 5 CV of buffer A supplemented with 300 mM imidazole. The precursor-chaperone complex was immediately subjected to dialysis against buffer A prior to concentrating on Amicon 30 kDa MWCO centrifugal filters (1000 g). Immediate removal of imidazol was particularly important for the preparation of the less stable Tim23_FL_-TIM9·10 complex. Complexes of Tim23_IMS_ with TIM8·13 or TIM9·10 were prepared by mixing two purified protein samples, and dialysis against buffer A. Formation of the precursor-chaperone complex was verified by Size-Exclusion Chromatography on Superdex 200 column. The resulting complex was further characterized by Size-Exclusion Chromatography coupled to Multi-Angle Light Scattering (SEC-MALS). TIM8·13 and TIM8·13-Tim23 were furthermore analyzed by analytical ultra-centrifugation (AUC). Both experiments were performed at 10°C in Buffer A. The amount of eluted complex was estimated from the protein concentration, measured absorbance of the sample at 280 nm and the sum of the molecular weights and extinction coefficients of the chaperone and the precursor protein.

### Competition assays

The first competition assay was performed by adding an equimolar mixture of TIM8·13 and TIM9·10 chaperones to the NiNTA bound precursor protein, Tim23_FL_ or Ggc1. After washing the column, precursor-chaperone complex was eluted in Buffer A supplemented with 300 mM imidazole. In the time dependent competition assay, complex of precursor protein and one of the chaperones (TIM8·13 or TIM9·10) was prepared before adding the equimolar concentration of the other chaperone at the time point 0. The reaction mixture was incubated at 30°C. After 0.5, 1 and 3 hours, aliquot of the reaction mixture was taken and (newly formed) precursor-chaperone complex was isolated on a NiNTA affinity column. Difference in the amount of specific chaperone, TIM8·13 or TIM9·10, bound to precursor protein was analysed by SDS-PAGE and liquid chromatography coupled to mass-spectrometry (LC ESI-TOF MS, 6210, Agilent Technologies, at the Mass Spectrometry platform, IBS Grenoble). Samples for the analysis by mass spectrometry were heat shocked for 15’ at 90°C, resulting in the dissociation and precipitation of the precursor protein, while the apo-chaperones were recovered in the supernatant after cooling the sample and centrifugation for 10’ at 39k g. As a reference, samples of precursor proteins, Tim23_FL_ and Ggc1, bound to individual chaperone, TIM8·13 or TIM9·10, were prepared and analysed in parallel. To be noted, preparation of the TIM8·13-Ggc1 complex, in quantity sufficient for the analysis, was unsuccessful. To calculate the difference in the amount of specific chaperone bound to precursor protein, normalized areas under the chromatography peaks corresponding to each Tim monomer were used.

### Isothermal Titration Calorimetry experiments

Calorimetric binding experiments of Tim23_IMS_ and TIM chaperones were performed using a MicroCal ITC200 instrument (GE Healthcare). Sixteen successive 2.5 µl aliquots of 1.15 mM Tim23_IMS_ were injected into a sample cell containing 55 µM TIM9·10 or TIM8·13. All ITC data were acquired in Buffer A at 20°C. Control experiments included titrating Tim23_IMS_ into the Buffer A. The enthalpy accompanying each injection was calculated by integrating the resultant exotherm, which corresponds to the released heat as a funcSEC-MALStion of ligand concentration added at each titration point. ITC data were analysed via the MicroCal Origin software using a single site binding model and nonlinear least squares fit of thermodynamic binding parameters (ΔH, K, and n). We also performed ITC experiments with the VDAC peptides; no efSEC-MALSfects could be detected, in line with a millimolar affinity, as already reported for the TIM9·10–cyclic-VDAC_257-279_ pepSEC-MALStide (23).

### Cross-linking of VDAC_257-279_

*In vitro* cross-linking of VDAC_257-279_ used precisely the protocol described in ref. (23) for TIM9·10. Briefly, 5 µM TIM8·13 was mixed with VDAC_257-279_ at 0, 25 or 50 µM, incubated 10 min on ice, and UV-illuminated (30 min, 4 °C).

### SEC-MALS experiments

SEC-MALS experiments were performed at the Biophysical platform (AUC-PAOL) in Grenoble. The experimental setup comprised a HPLC (Schimadzu, Kyoto, Japan) consisting of a degasser DGU-20AD, a LC-20AD pump, a autosampler SIL20-ACHT, a column oven XL-Therm (WynSep, Sainte Foy d’Aigrefeuille, France), a communication interface CBM-20A, a UV-Vis detector SPD-M20A, a static light scattering detector miniDawn Treos (Wyatt, Santa-Barbara, USA), a dynamic light scattering detector DynaPro NANOSTAR, a refractive index detector Optilab rEX. The samples were stored at 4 °C, and a volume of 20, 40, 50 or 90 µl was injected on a Superdex 200, equilibrated at 4 °C; the buffer was 50 mM Tris, 150 mM NaCl filtred at 0.1 µm, at a flow rate of 0.5 ml/min. Bovine serum albumine was used for calibration. Two independent sets of experiments conducted with two different batches of protein samples were highly similar.

### Analytical ultra-centrifugation

AUC experiments of TIM8·13 and TIM8·13-Tim23 were performed at 50000 rpm and 10 °C, on an analytical ultracentrifuge XLI, with a rotor Anti-60 and anti-50 (Beckman Coulter, Palo Alto, USA) and double-sector cells of optical path length 12 and 3 mm equipped of Sapphire windows (Nanolytics, Potsdam, DE). Acquisitions were made using absorbance at 250 and 280 nm wave length and interference optics. The reference is the buffer 50 mM Tris, 150 mM NaCl. The data were processed by Redate software v 1.0.1. The c(s) and Non Interacting Species (NIS) analysis was done with the SEDFIT software, version 15.01b and Gussi 1.2.0, and the Multiwavelenght analysis (MWA) with SEDPHAT software version 12.1b.

### NMR spectroscopy

All NMR experiments were performed on Bruker Avance-III spectrometers operating at 600, 700, 850 or 950 MHz ^1^H Larmor frequency. The samples were in the NMR buffer (50 mM NaCl, 50 mM Tris, pH 7.4) with 10% (v/v) D_2_O, unless stated differently. All multidimensional NMR data were analyzed with CCPN (version 2) (40). DOSY data were analyzed with in-house writSEC-MALSten python scripts. For calculating chemical-shift perturSEC-MALSbation data, the contribution of each different nuclei was weighted by the gyromagnetic ratios of the respective nuSEC-MALScleus: e.g. the combined ^1^H-^15^N CSP was calculated as 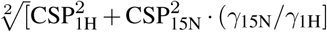, where the *γ* are the gyroSEC-MALSmagnetic ratios.

#### TIM8·13 and Tim23_IMS_ resonance assignments

For the resSEC-MALSonance assignment of TIM8·13, the following experiments were performed : 2D ^15^N-^1^H-BEST-TROSY HSQC, 3D BEST-TROSY HNCO, 3D BEST-TROSY HNcaCO, 3D BEST-TROSY HNCA, 3D BEST-TROSY HNcoCA, 3D BEST-TROSY HNcocaCB and 3D BEST-TROSY HNcaCB (41, 42) and a 3D ^15^N-NOESY HSQC. The experiments were performed with a 0.236 mM [^2^H,^15^N,^13^C]-labeled TIM8·13,at 308 K and 333K. The NMR resonance assignment of TIM9·10 was reported earlier (23). We collected BEST-TROSY HNCA, HNCO and HNcoCA experiments to assign Tim23_IMS_, aided by the previously reported assignSEC-MALSment (37).

#### VDAC titration experiments

Cyclic-hVDAC1_257-279_ peptide was synthesized and lyophilized as described elsewhere ((35)). The peptide was dissolved in DMSO, and the DMSO concentration was reduced to 10% by step-wise addition of NMR buffer (1:1 in each step). Chaperone, TIM9·10 or TIM8·13, in buffer A was added to yield a final DMSO conSEC-MALScentration of 6% and a chaperone concentration of 0.15 mM (TIM9·10) or 0.1 mM (TIM8·13). Combined ^15^N-^1^H chemSEC-MALSical shift-perturbation (CSP) was calculated from the chemSEC-MALSical shifts obtained from the ^15^N-^1^H HSQC spectra of the complex samples with molar ratio of 1:4 for TIM9·10:VDAC, and ratio of 1:5 for TIM8·13:VDAC, in comparison to the chemical shifts from the apo-chaperone spectrum. The NMR experiments were performed at 308K.

#### Tim23_IMS_ titration experiments

For each titration point indiSEC-MALSvidual samples were prepared by mixing two soluble proSEC-MALStein samples, and monitored using ^15^N-^1^HSEC-MALSBEST-TROSY HSQC experiments at 283K (for Tim23 observed experiSEC-MALSment) or at 308K (for chaperone observed experiments). Titration samples with 100 µM [^15^N]-labeled Tim23_IMS_ with the molar ratios for Tim23_IMS_:TIM8·13 from 1:0 to 1:4, and the molar ratios for Tim23_IMS_:TIM9·10 from 1:0 to 1:5, were used. For the chaperone observed experiments, used samples contained 200 µM [^2^H,^13^C,^15^N]-labeled TIM8·13 with molar ratios of Tim23_IMS_ 1:0 and 1:1, and 350 µM [^2^H,^13^C,^15^N]-labeled TIM9·10 with 1:0 and 1:3 molar ratios of Tim23_IMS_.

#### NMR experiments with the Tim23_FL_

Complexes of the chapSEC-MALSerones with the full-length Tim23 were prepared as indiSEC-MALScated above (Preparation of chaperone-preursor protein comSEC-MALSplexes). Peak positions (chemical shifts) of the amide backbone sites of TIM8·13, apo-and in the complex with Tim23_FL_, were obtained from the ^1^H-^15^N HSQC exSEC-MALSperiments at 308K, with 120 µM [^13^CH_3_-ILV]-TIM8·13SEC-MALSTim23_FL_ sample. Similarly, to calculate combined ^15^N-^1^H and ^13^C-^1^H CSPs, chemical shifts of the amide backbone and ILVA-^13^CH_3_ groups of TIM9·10, apo-and in the complex with Tim23_FL_, were obtained from the ^1^H-^15^N HSQC and ^1^H-^13^C HMQC experiments at 288K. Sample of the [^13^CH_3_-ILVA]-TIM9·10 with the Tim23_FL_ was at 140 µM concentration. For the CSP calculations with the complexes of [^2^H-^15^N]-labelled Tim23_FL_ and the chaperones (190 µM complex with TIM8·13,and 61 µM complex with TIM9·10), chemical shifts from ^1^H-^15^N HSQC experiments at 288K were used in comparison to the chemical shifts of the apo-Tim23_IMS_.

#### Diffusion ordered spectroscopy

Diffusion-ordered NMR spectroscopy (DOSY) experiments were performed at 288K and 600 MHz ^1^H Larmor frequency. Diffusion constants were derived from a series of one-dimensional ^1^H spectra either over the methyls region (methyl-selective DOSY experiments, for ^13^CH_3_-ILVA-labeled apo-and Tim23_FL_ bound TIM9·10) or over the amides region (for [^15^N]Tim23_FL_TIM9·10). Diffusion coefficients were obtained from fitting integrated 1D intensities as a function of the gradient strength at constant diffusion delay.

### Small-angle X-ray scattering data collection and analysis

SAXS data were collected at ESRF BM29beam line (43) with a Pilatus 1M detector (Dectris) at the distance of 2.872 m from the 1.8 mm diameter flow-through capillary. Data on TIM8·13 were collected in a batch mode. The X-ray energy was 12.5 keV and the accessible q-range 0.032 nm^*–*1^ to 4.9 nm^*–*1^. The incoming flux at the sample position was in the order of 1012 photons/s in 700×700 mm^2^. All images were automatically azimuthally averaged with pyFAI (44). SAXS data of pure TIM8·13 was collected at 1, 2.5 and 5 mg/mL using the BioSAXS sample changer (45). Ten frames of one second were collected for each concentration. Exposures with radiation damage were discarded, the remaining frames averaged and the background was subtracted by an online processing pipeline (46). Data from the three concentrations were merged following standard procedures to create an idealized scattering curve, using Primus from the ATSAS package (47). The pair distribution function p(r) was calculated using GNOM (48).

Online purification of the TIM8·13–Tim23(FL) complex using gelfiltration column (HiLoad 16/600 Superdex S200 PG) was performed with a high pressure liquid chromatography (HPLC) system (Shimadzu, France), as described in reference (49). The HPLC system was directly coupled to the flow-through capillary of SAXS exposure unit. The flow rate for all online experiments was 0.2 mL/min. Data collection was performed continuously throughout the chromatography run at a frame rate of 1 Hz.

### Molecular dynamics simulations and fitting of SAXS data

The inital model of Tim23 was built using I-TASSER (Iterative Threading ASSEmbly Refinement) (50) and QUARK webservers (51) which predicted a long unstructured N-terminal tail and four/five helical structures in the transmembrane domain. The structure of TIM9·10 hexamer built in our previous work (23) was used as the initial model of TIM9·10 chaperone, and as the template to build the model of TIM8·13 chaperone based on the sequence of yeast Tim8 and Tim13 (UniProt IDs: P57744 and P53299) by homology modeling with MODELLER (52). (Note that in the crystal structure of TIM8·13 (PDB-ID 3CJH), more than 75 residues are missing in each Tim8-Tim13 pair, thus requiring model building.) The disulfide bonds related to the twin CX_3_C motif were also kept in these models. The structures of the TIM8·13 hexamer and Tim23 were subsequently used to build the full structure of the TIM8·13-Tim23 complex by manually wrapping the helical structures of the transmembrane domain of Tim23 around the hydrophobic cleft of TIM8·13 which has been identified by NMR, and leaving the unstructured N-terminal of Tim23 as a floppy tail. The complex structure was further optimized by energy minimization and relaxation in 100 ns MD simulations using the simulation protocol as described in the following section. This model was used to generate the so-called ‘N-tail unbound’ ensemble in which the N-terminal half of Tim23 is free in solution. Based on the ‘N-tail unbound’ model of the TIM8·13-Tim23 complex, we further constructed the ‘N-tail bound’ model in which the N-terminal half of Tim23 is in contact with the upper part of TIM8·13. This was achieved by adding a restraint term in the force field using PLUMED plugin (53), *V*_*restraints*_, which is a half-harmonic potential of the form of *k*(*R* –*R*_0_)^2^ when R is larger than *R*_0_, and zero when R less than *R*_0_. Here R is the distance between the center of mass of Tim23Nter and the top part of the chaperone. We used *R*_0_ = 1*nm* and k = 400 *kJmol*^*–*1^. The ‘N-tail bound’ and unbound models for the TIM9·10-Tim23 complex were constructed by replacing TIM8·13 with TIM9·10 based on the corresponding TIM8·13-Tim23 models.

The TIM8·13-Tim23 complex in the ‘N-tail unbound’ conformation was placed into a periodic cubic box with sides of 17.5 nm solvated with TIP3P water molecules containing Na^+^ and Cl^*–*^ ions at 0.10 M, resulting in ∼700,000 atoms in total. To reduce the computational cost, the complex in ‘N-tail bound’ conformation was placed in a smaller cubic box with sides of 12.9 nm, resulting in ∼300,000 atoms in total. The systems of the TIM9·10-Tim23 complex have similar size as the TIM8·13-Tim23 systems in the corresponding states. The apo TIM8·13 chaperone was placed into a periodic cubic box with sides of 12.0 nm, containing ∼230,000 atoms.

The Amber ff99SB-disp force field (54) was used for all simulations. The temperature and pressure were kept constant at 300 K using the v-rescale thermostat and at 1.0 bar using the Parrinello-Rahman barostat with a 2 ps time coupling constant, respectively. Neighbor searching was performed every 10 steps. The PME algorithm was used for electrostatic interactions. A single cut-off of 1.0 nm was used for both the PME algorithm and Van der Waals interactions. A reciprocal grid of 96 × 96 x 96 cells was used with 4th order B-spline interpolation. The hydrogen mass repartitioning technique (55) was employed with a single LINCS iteration (expansion order 6) (56), allowing simulations to be performed with an integration time step of 4 fs. MD simulations was performed using Gromacs 2018 or 2019 (57).

A total of 4.25 µs trajectories were collected to sample the conformational space of the chaperone-Tim23 complexes in both ‘N-tail bound’ and ‘N-tail unbound’ states. Four *µs* trajectories were also collected to sample the ensemble of apo TIM8·13 chaperone. These sampled conformations were used for further ensemble refinement using the Bayasian Maximum Entropy (BME) method (38, 39) guided by experimental SAXS data as decribed in our previous work (23). The distribution of both states in principle could be identified from the force field but needs substantial sampling. Therefore instead of estimating the prior by large-scaled MD simulations, we assigned equal weight (50%) for both states by inputting equal number of conformations (5000) into the mixed ensemble so without bias to either state. By tuning the regularization parameter in the BME reweighting algorithm, we adjusted the conformational weights in variant degrees to improve the fitting with experimental SAXS data.

The hydrogen bond and salt bridge formation between the N-terminal tail of Tim23 (residue 1-100) and the top surface of the chaperones was analyzed by GetContacts scripts (https://getcontacts.github.io/), and visulized using Flareplot (https://gpcrviz.github.io/flareplot/). Protein structures were visualized with PyMOL (58) and VMD (59).

### Calculations of affinities and populations

#### Estimation of the population of ‘N-tail bound’ states from ITC-derived K_d_

We attempted to link the ITC-derived dissociation constant of the Tim23_IMS_ fragment to the populations of bound and unbound states in the Tim23_FL_-chaperone complexes, using a rationale akin to the one outlined earlier for binding of disordered proteins to two sub-sites (60). Briefly, we treat the N-terminal tail of Tim23 as a ligand, and the remaining bound complex as the target protein, then the relationship between the population of the bound state (P_bound_) and the binding affinity (K_d_) can be written as P_bound_/(1-P_bound_) = C_eff_/K_d_ where C_eff_ is the effective concentration of the disordered N-tail which was estimated to be between 0.2-3 mM from the MD simulations, resulting in the estimation of P_bound_ to be between 75% and 98%.

#### Estimation of the K_d_ ratio from competition assays

Determining dissociation constants of TIM chaperones to its insoluble client proteins is hampered by the impossibility to form the complexes by solution methods such as titration, as requires the pull-down method outlined in Fig. 1A. Nonetheless, the amount of TIM8·13-Tim23 and TIM9·10-Tim23 complexes obtained in the competition assays (Fig. 1) can provide an estimate of the relative affinities. The dissociation constants can be written from the concentrations as follows:

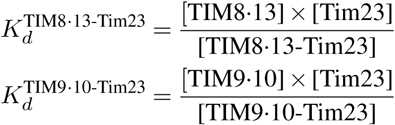

where [TIM8·13-Tim23] denotes the concentration of the formed chaperone-precursor complex, and [TIM8·13] and [Tim23] are the concentrations of free chaperone and precursor protein in solution. The latter is negligible, as no free precursor protein is eluted from the column (some aggregated precursor protein was removed from the equilibrium). Both chaperones have been applied at the same concentration *c*_0_ = [TIM8·13]+[TIM8·13-Tim23] = [TIM9·10]+[TIM9·10-Tim23] to the resin-bound precursor protein that was present at a concentration *b*_0_ = [Tim23]+[TIM9·10-Tim23]+[TIM8·13-Tim23]. Using the ratio of formed complex obtained in the competition assay, *r* = [TIM9·10-Tim23]/[TIM8·13-Tim23] leads to

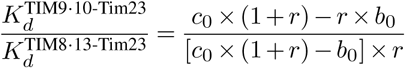

The experimental protocol does not allow to determine with precision the concentrations of precursor protein (*b*_0_) and each chaperone (*c*_0_), as the former is bound to a resin. As the chaperone was added in excess, and some of the precursor protein precipitated on the column, we can safely assume *c*_0_ ≤*b*_0_. With an experimentally found ratio of formed complexes of *r* = 5 and assuming that *c*_0_*/b*_0_ assumes the values of 1-5, the K_d_ ratio falls in the range of 1:25 to 1:6, i.e. ca. one order of magnitude.

##### Acknowledgements

We thank Aline Le Roy and Christine Ebel, and Luca Signor for excellent support with SEC-MALS experiments and mass spectrometry experiments, respectively. We thank Hubert Kalbacher (Univ. Tübingen) for synthesis of the VDAC peptides, and Karine Giandoreggio-Barranco for preparing Tim23 samples and Undina Guillerm for support with protein production. We are grateful to Bernhard Brutscher, Alicia Vallet and Adrien Favier for excellent NMR platform operation and management. We acknowledge insightful discussion with Nils Wiedemann and Caroline Lindau. This study was supported by the European Research Council (StG-2012-311318-ProtDyn2Function), and the Agence Nationale de la Recherche (ANR-18-CE92-0032). This study used the platforms (NMR, EM, isotope-labeling) of the Grenoble Instruct-ERIC Center (ISBG; UMS 3518 CNRS-CEA-UJF-EMBL) with support from FRISBI (ANR-10-INSB-05-02) and GRAL (ANR-10-LABX-49-01) within the Grenoble Partnership for Structural Biology (PSB). Y. W. and K. L.-L. were supported by the BRAINSTRUC structural biology initiative from the Lundbeck Foundation. We acknowledge access to computational resources from the Danish National Supercomputer for Life Sciences (Computerome) and the ROBUST Resource for Biomolecular Simulations (supported by the Novo Nordisk Foundation). This work was furthermore supported by the Deutsche Forschungsgemeinschaft (RA 1028/8-1,2 to D. R.)

## Author contributions

K. W. and P. S. initiated this project. I. S., A. H., O. D., B. B. and K. W. prepared protein samples. I.S. performed the pull-down and competition assays. I.S., K. W., B. B., O. D. and P. S. performed and analyzed NMR data. K. W. and M. B. performed SAXS experiments. Y. W. and K. L.-L. performed and analyzed MD simulations. T. J. and D. R. performed/analyzed cross-linking experiments. All authors contributed ideas and discussion. I.S., Y. W. and P.S. made the figures. P. S., I. S. and Y. W. wrote the manuscript with input from all authors.

## Data availability

The chemical shift assignments of TIM8·13 have been deposited in the BioMagResBank (www.bmrb.wisc.edu) under accession number 50213. All MD models and SAXS data have been deposited in the SASBDB (www.sasbdb.org) under accession number SASDH89 (TIM8·13-Tim23), SASXXX (TIM8·13), SASYYY (TIM9·10-Tim23) and SASDEF2 (TIM9·10 (23)). All chemical-shift perturbation data have been deposited on Mendeley data (http://dx.doi.org/10.17632/8cr8rvtddm.2).

---

**Fig. S1.**
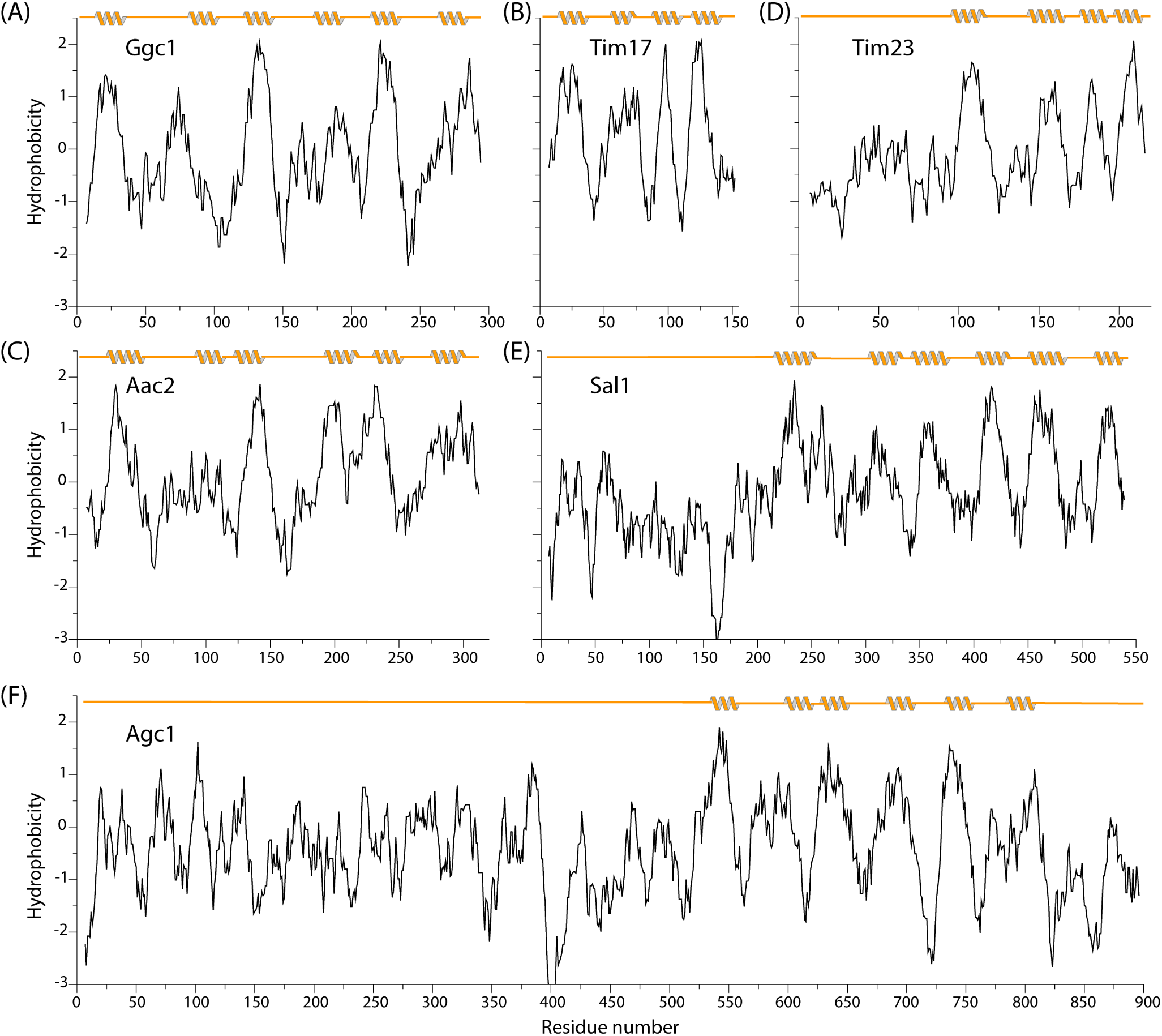
Kyte-Doolittle hydrophobicity of membrane precursor proteins known not to bind to TIM8·13 (Tim17, Ggc1, Aac2) and proteins known to bind (Glu-Asp carrier Agc1, Tim23 and, putatively, Sal1 - the human form of Sal1, Citrin, is known to depend on the human homolog of TIM8·13). The Kyte-Doolittle hydrophobicity (61) has been determined with the web server of ExPASy, using a window size of 13 and the standard linear weight variation model. Trans-membrane helices (but not other secondary structure elements) are indicated above each plot, as determined from either UniProt or a modelling with SwissModeller (for Sal1, using the structure of Aac2 as a template).

**Fig. S2.**
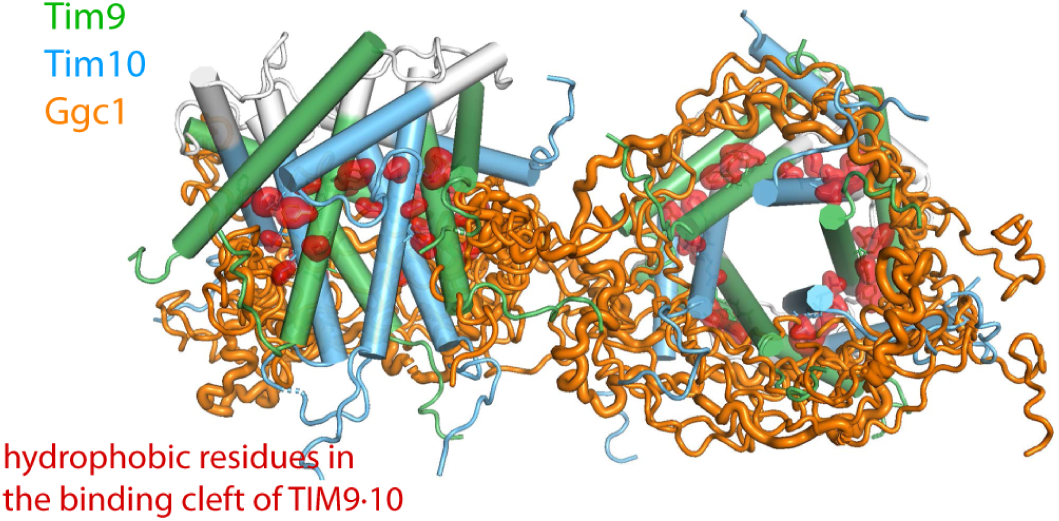
Ensemble representation of the structure of TIM9·10 (blue/green) holding full-length Ggc1 (orange), as reported in ref. (23). In contrast to the complexes presented in this study, the complex has a 2:1 stoichiometry, and essentially the entire precursor protein is located to the hydrophobic binding cleft of the chaperone.

**Fig. S3.**
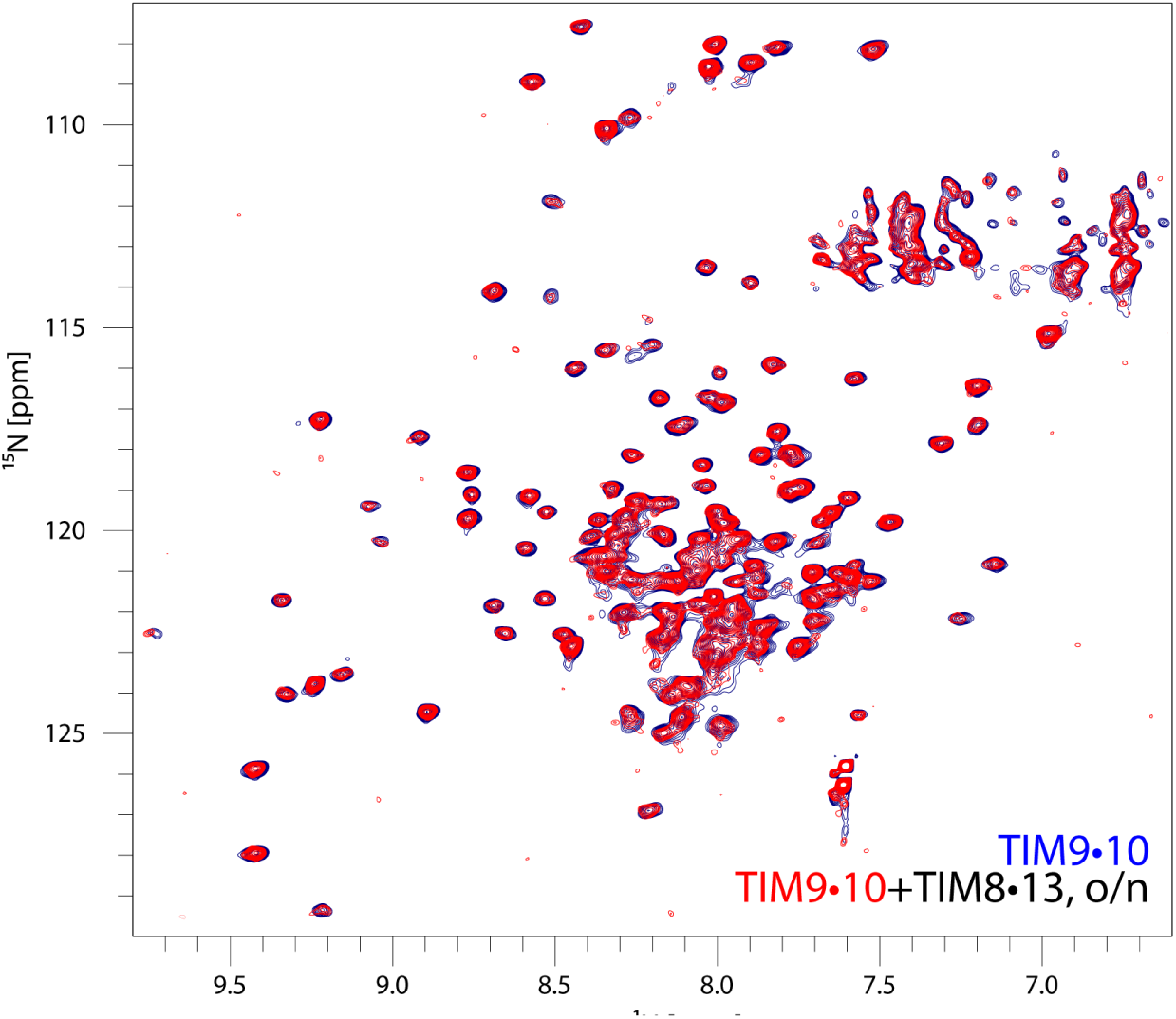
TIM8·13 and TIM9·10 do not form mixed complexes. NMR spectra of a ^2^ H,^15^ N,^13^ C labelled TIM9·10 sample (blue) and a mixture of this sample with unlabelled (NMR-invisible) TIM8·13 after overnight incubation. If TIM9·10 formed mixed oligomers with TIM8·13, the environment around each of the Tim9 and Tim10 subunits, as they would be surrounded by Tim8 or Tim13 subunits, would be different. Thus, the spectrum of TIM9·10 would feature additional peaks corresponding to those alternate environments. The spectrum after over-night incubation does not show any additional peaks and features, only the one set of peaks corresponding to the hexameric TIM9·10. Therefore, this data demonstrates that TIM9·10 does not form mixed hetero-oligomers with TIM8·13.

**Fig. S4.**
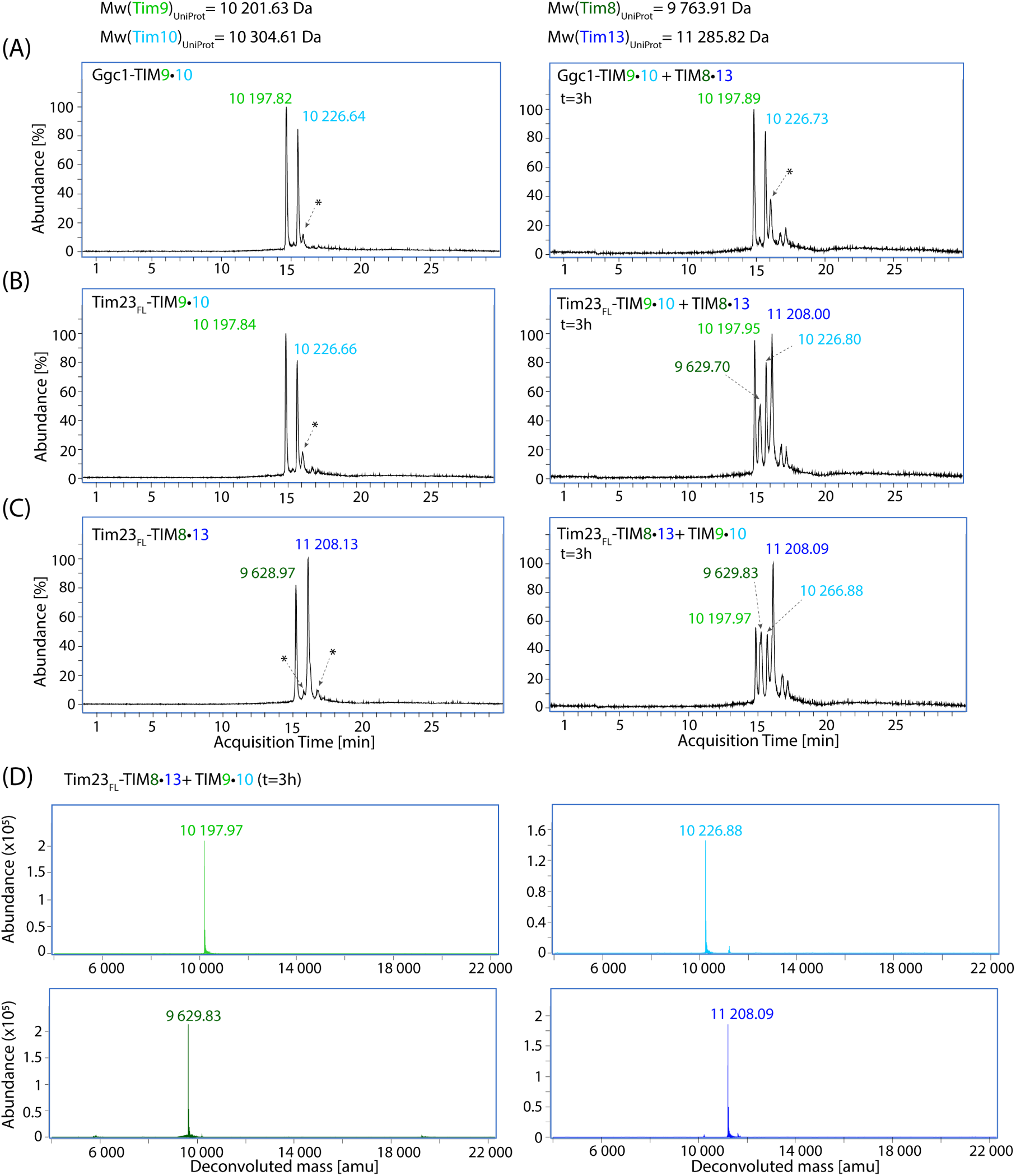
Liquid chromatography coupled with electrospray ionization, time-of-flight mass spectrometry (LC/ESI–TOF MS) analysis used to read out the difference in the amount of specific chaperone, TIM8·13 and TIM9·10, bound to precursor protein. (A) Mass chromatograms of the control sample, complex of TIM9·10 chaperone bound to Ggc1 (left), and the competition reaction three hours after adding TIM8·13 to the pre-formed Ggc1–TIM9·10 complex (right). Deconvoluted mass values are indicated next to corresponding peaks. Mass obtained for the unspecific peak, indicated with asterisk, was 10 226.53 (left) and 10 226.53 amu (right). (B) Same as in panel A, with the Tim23_FL_ as a substrate precursor protein. Mass of the unspecific peak in the left chromatogram was 10 198.69 amu. (C) Mass chromatograms of the TIM8·13 chaperone bound to Tim23_FL_ (left) as a control, and the competition reaction three hours upon adding TIM9·10 to the pre-formed Tim23_FL_–TIM8·13 complex (right). In the left chromatogram, obtained mass from the unspecific peaks (impurities) was 10 207.94 (left asterisk) and 11 208.27 amu (right asterisk). (D) Deconvoluted mass spectra for each of the chromatography peaks from the chromatogram shown in panel C, right. Reaction mixtures were heat-shocked before the analysis, resulting in the presence of the chaperone only in the analysed sample (see methods).

**Fig. S5.**
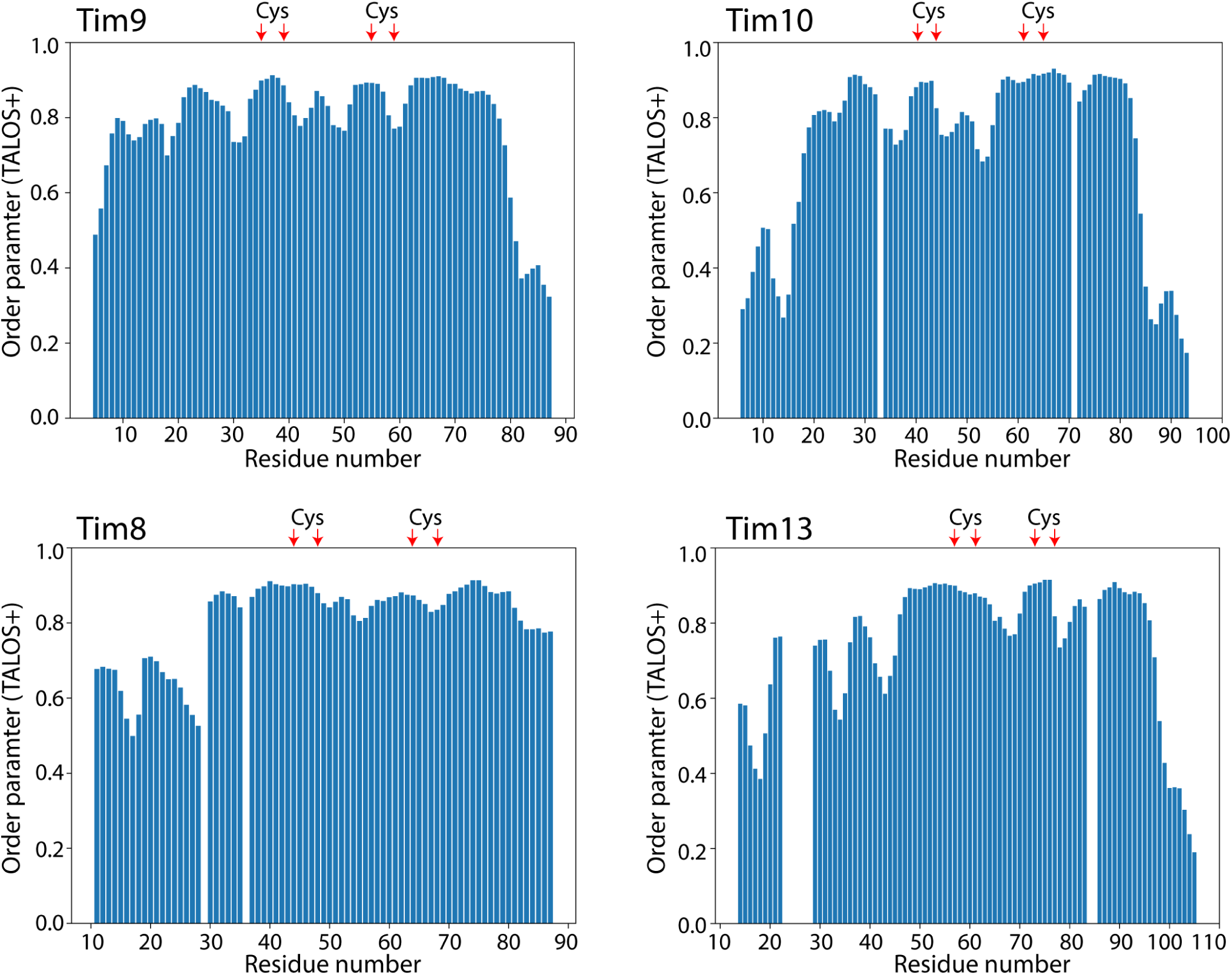
Local order parameters in TIM8·13 and TIM9·10, derived from assigned backbone chemical shifts and the TALOS-N (62) software. This data shows that the core of the chaperones is rather rigid, and the tentacles become increasingly flexible towards the N-and C-termini.

**Fig. S6.**
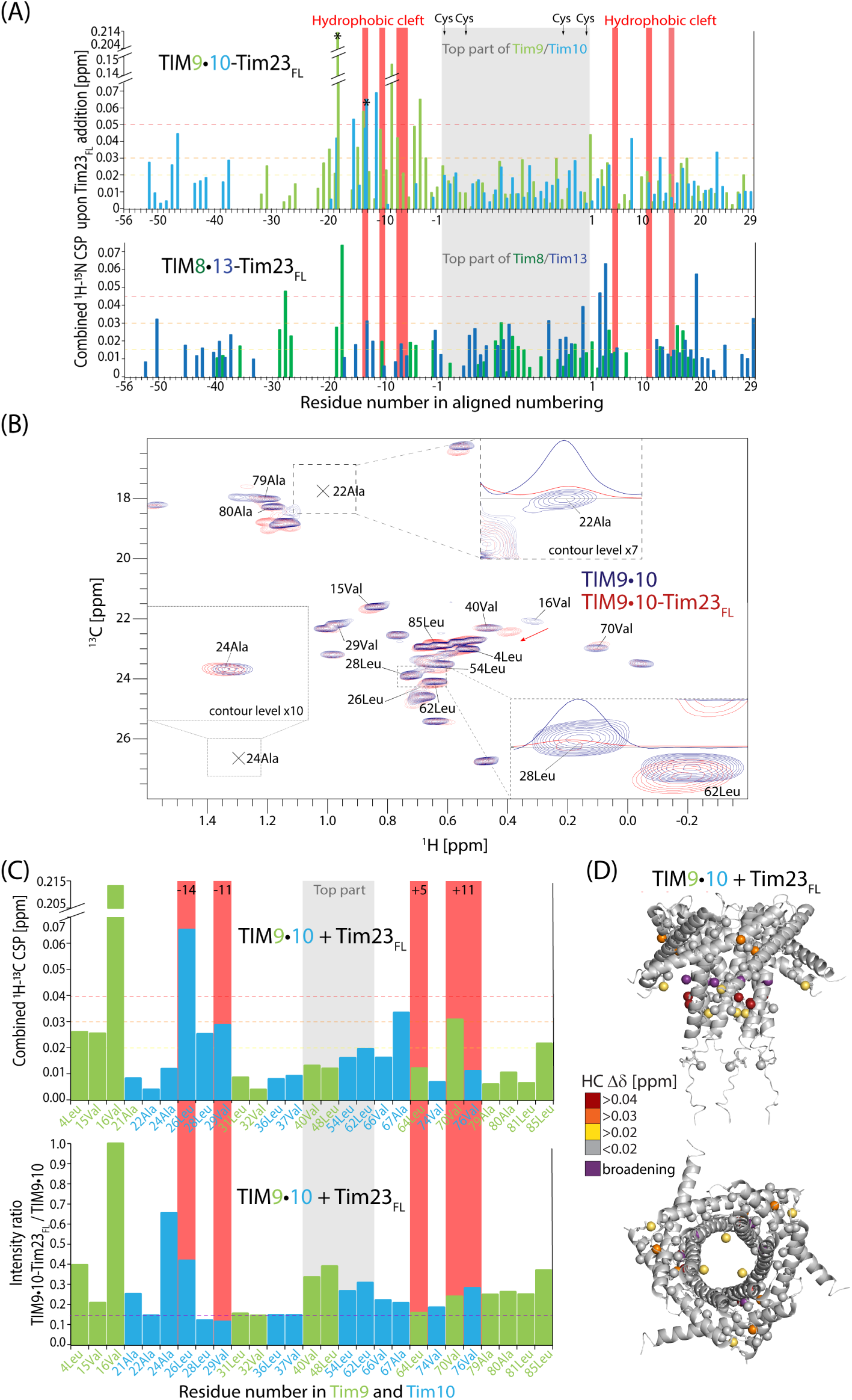
Full-length Tim23 interactions with TIM9·10 and TIM8·13 chaperones. (A) Chemical-shift perturbations observed upon addition of the Tim23_FL_ to TIM9·10 (top) and TIM8·13 (bottom). Comparison of CSPs on TIM8·13 induced by Tim23_FL_ and VDAC_257-279_ (Fig. 2C), shows that for binding of full-length Tim23, besides conserved hydrophobic binding site, TIM8·13 employs its hydrophilic top region where 3 fold increase in CSPs can be seen (mapped on the structure in Fig. 4C). Asterisk marks residues 16Val and 26Leu which showed biggest methyl CSP upon adding Tim23_FL_. (B) Methyl spectra overlay of the ILVA-labeled apo-TIM9·10 chaperone sample (blue) and ILVA-TIM9·10 in complex with the full-length Tim23 (red). In the zoomed-in regions, showing resides 22Ala and 28Leu, difference in the peak intensity is shown with the 1D traces (hydrogen spectra) at the corresponding carbon frequency of the selected peak. The biggest methyl CSP for the residue 16Val is indicated with an red arrow. (C) Methyl-detected CSP of the ILVA-labeled TIM9·10 upon addition of the full-length Tim23 (top), and the intensity ratio of the same residues (bottom). (D) Mapping of Tim23_FL_-induced methyl CSPs and biggest peak-broadening on TIM9·10, showing absence of interaction with the top part of TIM9·10.

**Fig. S7.**
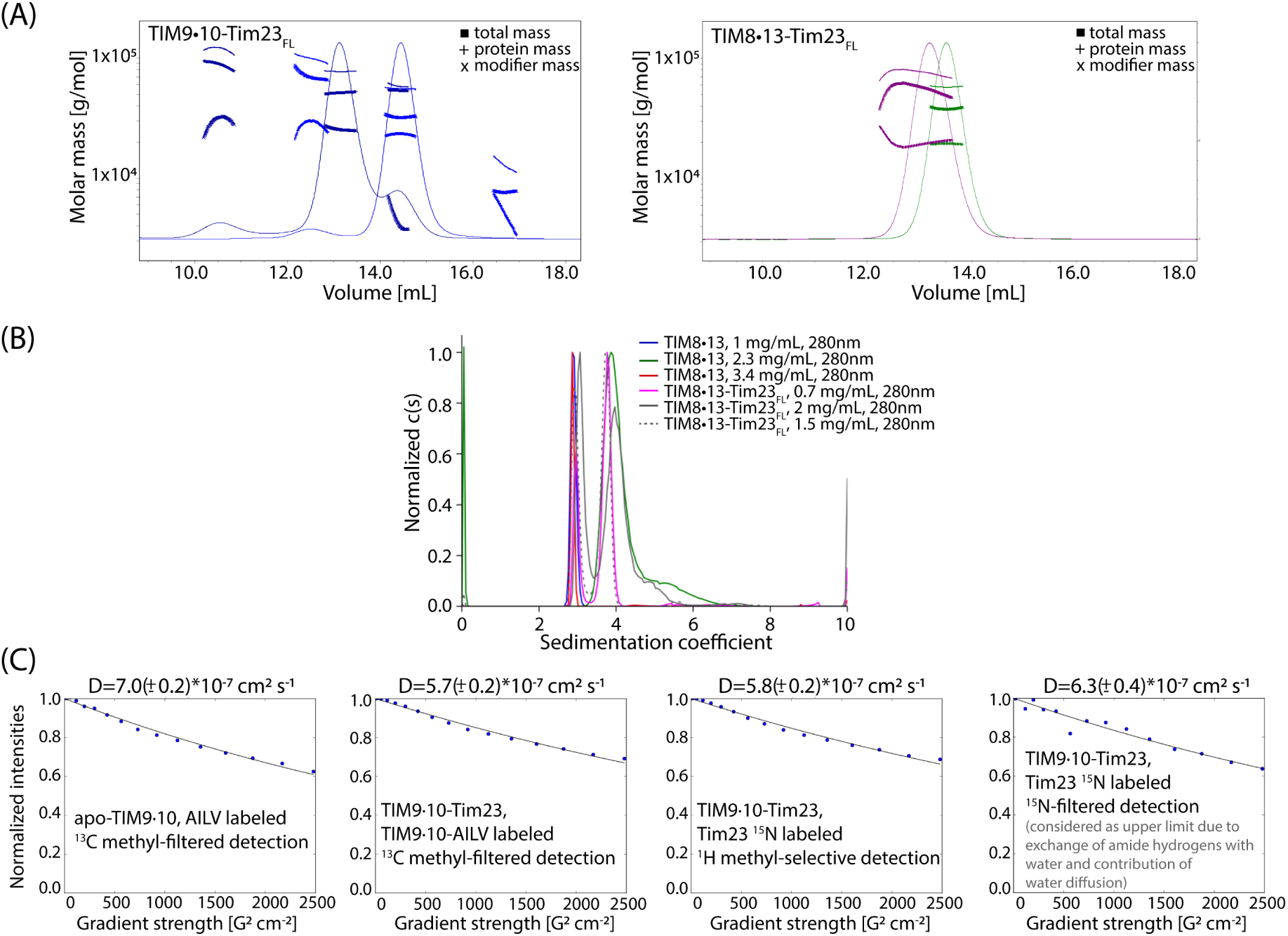
Experimental characterization of the size of TIM chaperones and their precursor protein complexes. (A) Size-exclusion-chromatography coupled to multi-angle light scattering (SEC-MALS) of TIM9·10–Tim23 and TIM8·13–Tim23. For TIM9·10 the dominant contribution (> 85 %) corresponds to 56 kDa, or ca. 5.5 subunits; for TIM9·10–Tim23, the major contribution (71 %) corresponds to a mass of 77.5 kDa, in reasonable agreement with the expected 23 kDa increase from Tim23-binding, with a clearly visible ‘shoulder’ at the position of the apo-TIM9·10. For TIM8·13, a peak corresponding to ca. 86 % of the signal is at a mass of 57.9 kDa; in TIM8·13–Tim23, the peak is at 74.2 kDa, in reasonable agreement with the expected 23 kDa increase. A control experiment with BSA yielded an observed mass of 62 kDa (theoretical: 66 kDa). (B) Analytical ultra-centrifugation of TIM8·13 and TIM8·13-Tim23 complexes. For the TIM8·13 we observe a main contribution, 98 +/- 1% of the total signal, at 2.89 +/- 0.01 S (s20w = 3.97 +/- 0.01S). The Non Interacting Species analysis gives M_w_ = 57 +/- 5 kDa. For the TIM8·13-Tim23 sample we observe a main contribution (67 +/- 2% of the total signal) at 3.76S (s20w = 5.17S), at 0.7 mg/mL sample concentration, slightly shifted to s20w = 5.67S, at 2 mg/mL. The Non Interacting Species analysis results in a M_w_ = 70 +/- 5 kDa, similar to SEC-MALS. A further contribution is detected at 3 +/- 0.05S (s20w = 4.1 +/- 0.1S), for 33 +/- 5 % of the total signal. The Non Interacting Species analysis gives M_w_ = 50 +/- 3 kDa, could be imprecise. This contribution superimposes to the main one of TIM8·13 alone. (C) NMR diffusion-ordered spectroscopy curves (DOSY), obtained as integrals of one-dimensional ^1^ H spectra over either methyls (first three) or amides (fourth panel) as a function of the gradient strength. The four panels correspond to three different samples, ^13^ CH_3_-ILVA-labeled apo-TIM9·10 (first panel), TIM9·10-ILVA methyl labeled TIM9·10- Tim23_FL_ complex (second panel) and Tim23_FL_-^15^ N labeled TIM9·10-Tim23_FL_ complex (last two panels). The first two used a ^13^ C-filtered experiment, the third one uses a ^1^ H methyl-frequency selective scheme and the fourth uses a ^15^ N-filtered version. The latter thus detects amide protons; as these are water exchangeable, in particular in disordered proteins, the observed diffusion coefficient is likely over-estimated, as it contains contributions from the diffusion coefficient of water; thus, this value is to be seen as an upper limit. The diffusion coefficient for a spherical particle scales with the cubic root of the molecular weight, which would lead to an expected change in the diffusion coefficient of (71.1/94.1)^1/3^, i.e. ca. 10 % lower in the complex compared to the apo state; additional domain motions, such as the flexibility of the Tim23 tail, decreases the diffusion coefficient, i.e. would lead to a larger relative change. This expected 10-15 % effect on diffusion coefficients is in good agreement with the experimental effect (11-20 %). A TIM9·10:Tim23 stoichiometry, as found for TIM9·10–Ggc1 would lead to an expected DOSY effect of 32 % (considering only mass), thus in worse agreement with the data.

**Fig. S8.**
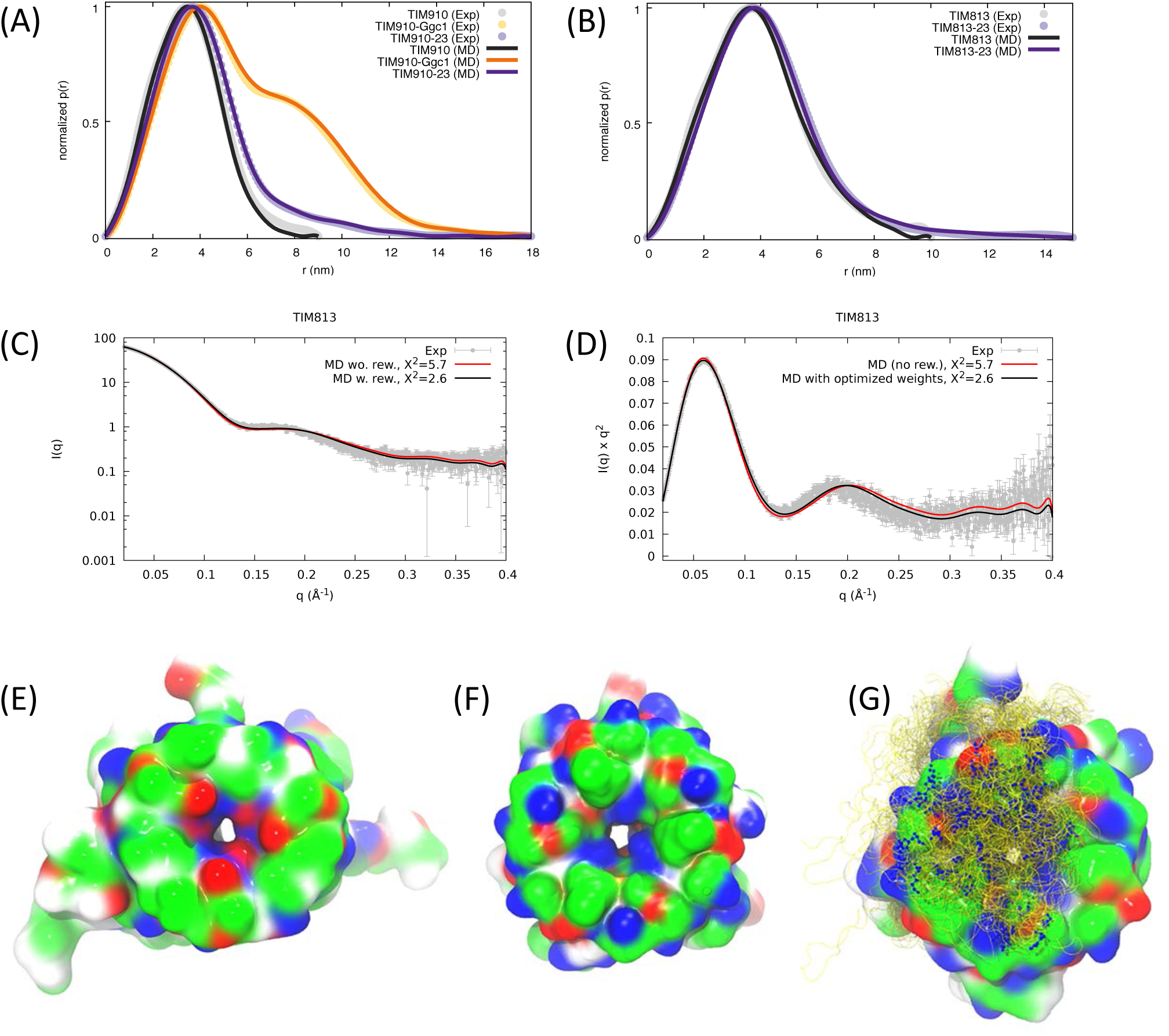
(A) P(r) distributions of TIM9·10 in the apo form and in complex with Tim23 or Ggc1. (B) P(r) distributions of TIM8·13 in the apo form and in complex with Tim23. (C) SAXS curves of TIM8·13 ensemble. (D) Kratky plot of SAXS curves of TIM8·13 ensemble. (E) Top view of a snapshot of MD simulations of apo TIM8·13. (F) Top view of a snapshot of MD simulations of apo TIM9·10. (G) Ensemble view of the N-tail-bound state of TIM9·10–Tim23.

